# Progressive Loosening of a Dual Autoinhibitory Interface Activates PP2A-B56δ

**DOI:** 10.64898/2026.08.19.745767

**Authors:** Michael Shea O’Connor, Cheng-Guo Wu, Yichong Lao, Yongna Xing, Xuhui Huang

## Abstract

Protein phosphatase 2A containing the B56δ regulatory subunit (PP2A-B56δ) is a critical signaling enzyme whose dysregulation is associated with cancer, neurodegenerative disorders, and Jordan’s syndrome, a severe intellectual disability disorder caused by mutations in B56δ. Unlike other PP2A holoenzymes, PP2A-B56δ is regulated through a unique dual autoinhibition mechanism in which the N- and C-arms occlude the catalytic site while a substrate-mimicking short linear motif (SLiM) blocks the substrate-binding pocket. Although disease-associated mutations have been shown to alter enzyme activity, the molecular mechanism underlying activation of PP2A-B56δ and the effects of pathogenic mutations remain poorly understood. Here, we combined cryo-electron microscopy (cryo-EM), enhanced-sampling molecular dynamics (MD) simulations, Markov state model (MSM) construction, and transition-state analysis using Transition State identification via Dispersion and vAriational principle Regularized neural networks (TS-DAR) to characterize the conformational landscape of the disease variant E198K. Our cryo-EM analysis identified two distinct structures of E198K: an inactive closed-form with the N/C-arms resolved and an active loose-form in which the N/C-arms become highly flexible and could not be fully resolved. These structures therefore established that activation is governed by conformational changes of the N/C-arms but did not reveal the underlying mechanism. Starting from the inactive closed-form, we generated over 1,600 trajectories with an average length of 1,260 ns combined for E198K and wild-type (WT) PP2A-B56δ. TS-DAR identified four metastable states and two major activation pathways connecting inactive and active conformations. We found that activation occurs through progressive loosening of the N/C-arm interface while maintaining the overall holoenzyme architecture, rather than a complete opening of the interface. This mechanism exposes both the catalytic site and substrate-binding pocket. Comparison of E198K and WT revealed that the disease-associated mutation shifts the conformational equilibrium toward active states while leaving the transition-state ensemble largely unchanged. Mechanistically, E198K disrupts a salt-bridge network and weakens interactions between the internal loop and the C-arm that normally stabilize active-site occlusion. The resulting increase in C-arm mobility promotes active-site exposure and explains the elevated catalytic activity of the mutant. Together, these findings establish a previously uncharacterized activation mechanism for PP2A-B56δ and provide an atomic-level explanation for how the pathogenic E198K mutation allosterically promotes holoenzyme activation.

## Introduction

PP2A is a major serine/threonine phosphatase that dephosphorylates protein substrates to regulate key signal transduction pathways. This family of enzymes is involved in crucial cellular processes such as cell cycle progression, growth, cell death and survival, and DNA damage response.^1–3^ The functional PP2A holoenzyme is a heterotrimer consisting of a common scaffold subunit A, a catalytic subunit C, and a variable regulatory subunit B, responsible for substrate recognition. The B56 family represents one of the four major classes of regulatory subunits, which includes the B56δ isoform.^4,5^ Mutations within B56δ regulatory subunit are implicated in human pathogenesis, including cancer,^6–10^ early-onset Parkinsonism,^11–13^ and intellectual disability. Notably, a set of 20 distinct *de novo* mutations in B56δ has been found in patients diagnosed with a range of intellectual disability symptoms, collectively named Jordan’s syndrome.^14–16^ Therefore, determining the molecular mechanisms underlying these B56δ mutations is essential to understanding PP2A-mediated disease.

Unlike other B56 family members, B56δ has a novel dual autoinhibition mechanism that remains largely elusive.^17^ This mechanism is mediated by the subunit’s uniquely long N- and C-terminal arms, which structurally block the active site of the catalytic subunit.^17^ Furthermore, the C-terminal arm features a tail that spans into the allosteric substrate-binding pocket. While B56 family members typically recognize substrates via short linear motifs (SLiMs) containing an LxxIxE sequence,^18–22^ the B56δ C-arm tail contains an intrinsic pseudo-substrate SLiM (the “C-arm SLiM”) that mimics these substrates. By competitively occluding the SLiM-binding pocket, this intrinsic motif acts in tandem with the physical blocking of the active site by the N/C-arms, establishing PP2A-B56δ’s distinctive dual autoinhibition.^17^

Previous studies have found that several of the Jordan’s syndrome mutations are located along the dynamic interface created by the N/C-arms.^17^ The presence of these mutations along this interface suggest they perturb the autoinhibition mechanism which results in a dysregulated PP2A-B56δ leading to detrimental diseases. Interestingly, these mutations are not directly involved in the catalytic mechanism or part of the substrate-binding site, yet they have been found to impair both catalysis and substrate binding.^17,23^ Therefore, the mutations must act allosterically in order to affect the dual autoinhibition mechanism. The most common and severe mutation from Jordan’s syndrome is E198K which results in the worse set of symptoms.^15,16,24,25^ Understanding this mutation’s effect on the autoinhibition of PP2A-B56δ is key to determining the disease-causing mechanism.

To elucidate the activation mechanism, one can use all-atom MD simulations to obtain atomic level dynamics of the dual autoinhibition mechanism of PP2A-B56δ. However, the timescale of this activation is expected to be beyond the capabilities of standard MD based on previous studies not successfully sampling N/C-arm loosening.^17,23^ Using MSMs, we can bridge the timescale gap by integrating hundreds of short MD simulations and discretizing the conformational landscape into metastable states.^26–35^ The resulting MSM can be used to predict the long timescales between these states.^36–46^ In addition to understanding the timescales of certain conformational changes, the identification of transition states between metastable states is important for understanding the structural transitions. Previous methods were only capable of identifying transition states for pairs of states individually.^47–49^ A newly developed method from our group called TS-DAR can automatically identify all candidate transition-state ensembles across free energy barriers between multiple metastable states.^50^ TS-DAR uses deep learning to detect transition-state candidate ensembles from out-of-distribution (OOD) conformations within a hyperspherical latent space. By optimizing kinetic compactness and inter-state dispersion on the hypersphere, the transition-state candidates can be easily identified by its OOD score and position on the latent space hypersphere. The transition-state conformations can be further validated by performing additional short MD simulations from candidates and show the equal probabilities of relaxing toward the adjacent metastable states. Mutations can affect the landscape by altering the free energy barrier between two basins or effecting the relative stability of these metastable states. Understanding a mutagenic effect on a transition state between two energy basins can provide valuable information on how the mutation effects the protein free energy landscape. Therefore, an efficient transition-state analysis is an effective tool for determining differences between WT and mutational variants.

Our study found that, contrary to the intuitive model of activation through complete opening of the N/C-arm interface, the activation occurs through progressive loosening of the interface while maintaining the overall architecture of the holoenzyme. Here, we combined cryo-EM, extensive all-atom MD simulations, TS-DAR analysis, and MSMs to elucidate this activation mechanism of the PP2A-B56δ disease-associated E198K variant. We resolved two cryo-EM structures of the E198K variant in an active loose-form conformation and in an inactive closed-form conformation. Interestingly, the loose-form conformation of E198K had a higher population compared to previously determined PP2A-B56δ E197K variant structures, consistent with increased enzymatic activity for this E198K mutation.^17^ However, the core enzyme remains nearly identical between the loose and closed structures, while the N/C-arms that regulate the dual autoinhibition become highly flexible and are largely unresolved in the loose-form reconstruction. Although the cryo-EM structures identify the inactive and loose conformations, they do not reveal how rearrangements of the N/C-arms drive activation of the holoenzyme. To reveal how the N/C-arms drive activation, we used MD simulations. Starting from the inactive closed-form structure of the E198K PP2A-B56δ holoenzyme, we generated over 1,600 trajectories with an average length of 1,260 ns across the E198K and WT systems. Our analysis identified four metastable states spanning the transition from a closed inactive state, in which the N/C-arms occlude the active site and the C-arm SLiM blocks the substrate-binding pocket, to an active state characterized by exposure of both regions. The activation process is initiated by displacement of the C-arm SLiM from the substrate-binding pocket and followed by sequential rearrangements of the N- and C-arms that expose the catalytic site. Furthermore, we show that the E198K mutation promotes these activating conformational changes by weakening interactions between the internal loop and the C-arm that normally stabilize the inactive state. Together, these findings establish a previously unrecognized activation mechanism for PP2A-B56δ and provide an atomic-level explanation for the hyperactivity of the E198K disease variant.

## Results and Discussion

### Two distinct cryo-EM structures of PP2A-B56δ E198K determined with greater loose-form population

We determined two cryo-EM structures of the disease-associated E198K variant of PP2A-B56δ. One structure is in a closed conformation with a resolution of 3.20 Å (Fig. 1E) and the other structure is in a loose conformation with a resolution of 2.93 Å (Fig. 1F) (see Methods Sections B-D for details). Based on the 3D variability analysis (3DVA)^51^ (Fig. S1), the ratio of loose/closed forms was found to be higher for E198K compared to the previously published structure of the E197K variant, which has more mild clinical severity.^14–16^ Although we failed to obtain the WT structure, previous binding assays showed that while E197K exhibits substrate SLiM binding similar to WT, the E198K variant displays a two-fold increase in binding, indicating it as a hyperactive variant.^17^ The greater ratio of loose form for E198K compared to the mild variant E197K is consistent with the increased substrate SLiM binding of E198K (Fig. S1).

**Figure 1.**
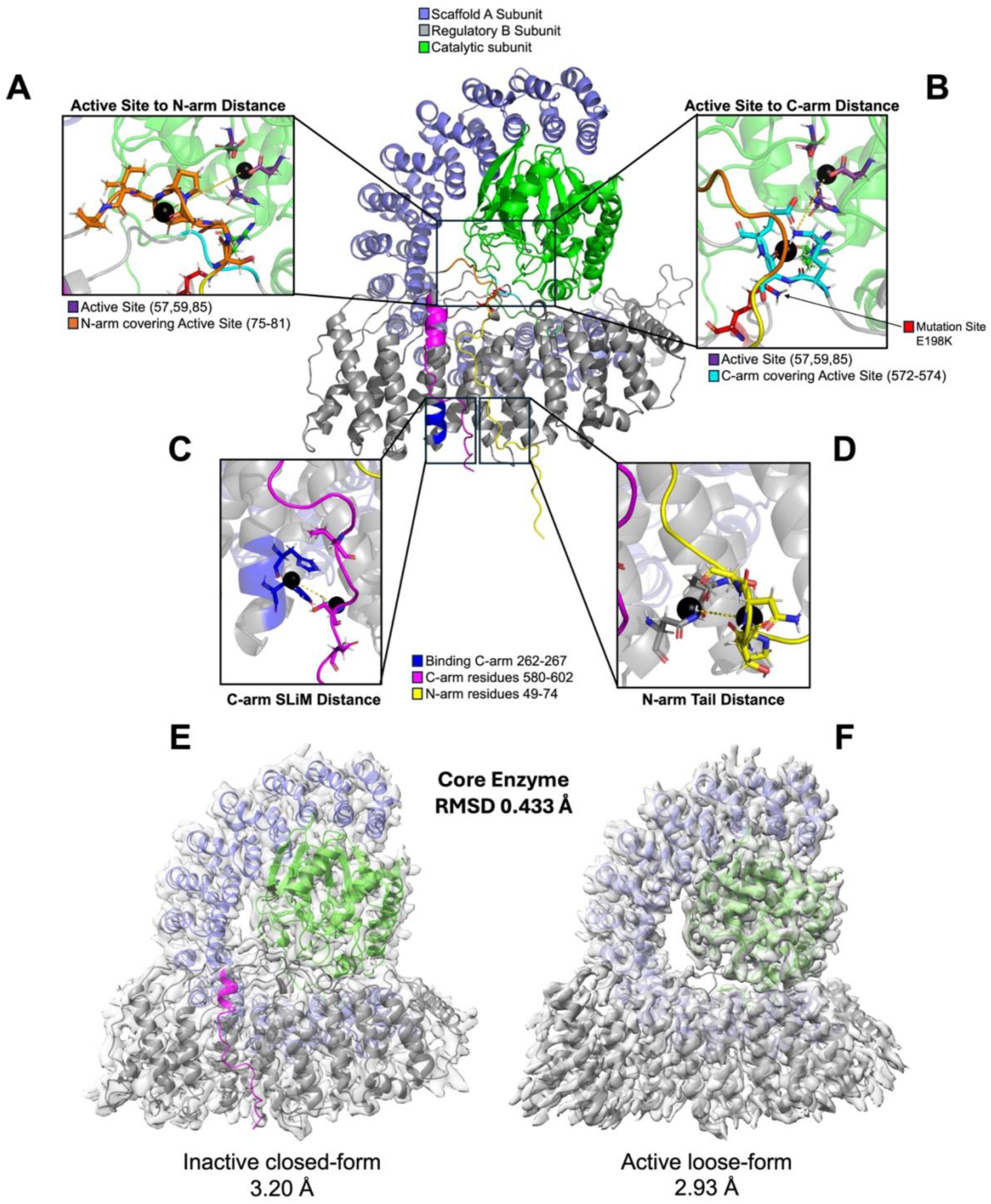
Cryo-EM structures of the E198K PP2A-B56δ holoenzyme and key distances to characterize conformational changes. The main panel shows the filled-in structural model of the PP2A-B56δ E198K mutant taken from the closed-form cryo-EM structure and used to initialize the all-atom MD simulations. **(A)** “Active Site to N-arm” distance is the center of mass (COM) distance between catalytic site residues D57, H59, D85 and regulatory subunit N-arm residues 75-81. This distance shows partial opening of the active site. **(B)** “Active Site to C-arm” distance is the COM distance between catalytic site residues and regulatory subunit C-arm residues 572-574 which shows partial opening of the active site. Together, the catalytic-site distances show the complete opening of the active site. **(C)** “C-arm SLiM” distance is the COM distance between the SLiM binding residues H263, Y266 and the C-arm SLiM residues L595, S598, E600. This distance measured the opening of the SLiM binding pocket. **(D)** “N-arm Tail” distance is the COM distance between regulatory subunit residues N309, Q349 to N-arm residues 59-62 which shows the flexibility of the N-arm tail. **(E)** Cryo-EM density and structure of inactive closed-form with a resolution of 3.20 Å. **(F)** Cryo-EM density and structure of active loose-form with a resolution of 2.93 Å. The core enzymes of each form are identical with an RMSD of 0.433 Å.

The core enzymes of the loose and closed forms are identical with an RMSD of 0.433 Å (Fig. 1E and F). However, similar to the reported structures of the E197K variant, the long N/C-arms of B56δ in the loose conformation do not interact with the holoenzyme core and remain structurally disordered.^17^ In this state, both the substrate-binding pocket and the catalytic active site are fully exposed. In contrast, the arms in the closed-form structure form extensive interactions with the core, allowing them to be fully resolved. Because the enzyme core remains similar between both states, the conformational switch is dictated by the structural state of these terminal arms. Therefore, we ran extensive MD simulations to sample the activation process of the holoenzyme, initialized from the closed-form structure.

In the closed-form structure, the substrate SLiM-binding pocket is blocked by the C-arm SLiM, while portions of the regulatory subunit N- and C-arms occlude the catalytic site, consistent with the closed-state structure of E197K variant.^17^ Because the N/C-arms occlude both the catalytic site and substrate-binding pocket in the closed form, we refer to the closed-form structure as the inactive state and the loose-form structure as the active state. In both structures, the E198K mutation site is located within an internal loop of the regulatory subunit spanning residues 190 – 200. Prior studies have shown that the E198K variant alters substrate binding and catalytic activity,^17,23^ this loop is not directly involved in either catalysis or substrate recognition, suggesting that the mutation exerts its effects through an allosteric mechanism.^23^ The internal loop residues 192-195 had lower local resolution in the loose form compared to the closed form indicating greater flexibility in the loose-form structure. Missing segments of the regulatory C-arm of the closed form, resulting from structural flexibility, were modeled using MODELLER^52^ (see Methods Section E for details). The resulting complete closed-form structure (Fig. 1) served as the starting point for all-atom MD simulations.

From the closed-form cryo-EM structure, we identified four key distances that describe the conformational state of the PP2A-B56δ holoenzyme and are used throughout this study. First, the “Active Site to N-arm” distance (Fig. 1A), measures the separation between active-site residues and N-arm residues that occlude the catalytic site, providing a metric for active-site accessibility based on the N-arm. Second, the “Active Site to C-arm” distance (Fig. 1B), quantifies the separation between the active site and the C-arm residues that cover it, providing a complementary measure of active-site openness. To characterize accessibility of the substrate-binding pocket, we define the “C-arm SLiM” distance (Fig. 1C) as the separation between residues in the regulatory subunit substrate-binding pocket and residues within the C-arm SLiM mimic. Finally, the “N-arm Tail” distance (Fig. 1D) measures the separation between the regulatory subunit core and the distal tail of the N-arm, providing a measure of N-arm flexibility. Together, these four distances capture the major conformational changes observed in the PP2A-B56δ holoenzyme.

### TS-DAR learns E198K dynamics and identifies four metastable states used for MSM construction

To sample the activation mechanism of the holoenzyme, we performed all-atom MD simulations starting from the closed-form E198K cryo-EM structure with the N/C-arms resolved. A previous study using unbiased all-atom MD was unable to sample opening of the active site and substrate-binding pockets due to the large timescale of these conformational changes.^23^ Here, we employed an enhanced sampling method, replica exchange with solute tempering (REST2),^53–55^ which scales the Hamiltonian potential of a selected region, called the “hot region”, of the protein across multiple replicas (see Methods Section F for details). Scaling the interactions within the “hot region” allows this region to move more freely, increasing its effective temperature and enabling greater conformational sampling. From the REST2 simulations, we projected each frame onto the “Active Site to C-arm” and “C-arm SLiM” distances and performed K-means clustering.^56^ This clustering yielded 176 representative structures spanning diverse conformations, which were subsequently used to seed extensive unbiased all-atom MD simulations on Folding@Home^57^ (see Methods Section G for details). The same procedure was applied to the WT system by reverting the E198K mutation in the initial cryo-EM structure. The total aggregate simulation time collected was 1.04 ms for E198K and 1.17 ms for WT (Fig. S2).

TS-DAR identified four macrostates for the E198K holoenzyme by learning the dynamic landscape from the MD simulations (Fig. 2A). Because the simulations were initiated from the E198K cryo-EM structure, we trained a four-state TS-DAR model using the extensive unbiased E198K trajectories. The overall machine learning architecture of TS-DAR is shown in Figure 2B. Training was performed on the time series of pairwise distances between selected key residues and filtered using Spectral Accelerated Sequential Incoherent Selection (Spectral-oASIS)^58^ (see Methods Section H for details). The TS-DAR model separated the conformational ensemble into four metastable states, each characterized by distinct structural features, and identified ensembles of transition-state candidates between specific macrostates using a three-dimensional hyperspherical embedding (Fig. 2C). Inspection of representative structures from each metastable state allowed us to assign biologically intuitive labels to the states: closed inactive (49.9%), intermediate 1 (12.2%), intermediate 2 (17.3%), and loose active (20.6%) (Fig. 2A). The closed-form cryo-EM structure corresponds to the closed inactive macrostate, whereas the loose-form cryo-EM reconstruction likely represents a heterogeneous ensemble spanning the intermediate 1, intermediate 2, and the loose active states because the increased flexibility of the N/C-arms prevents these conformations from being resolved as separate cryo-EM classes.

**Figure 2.**
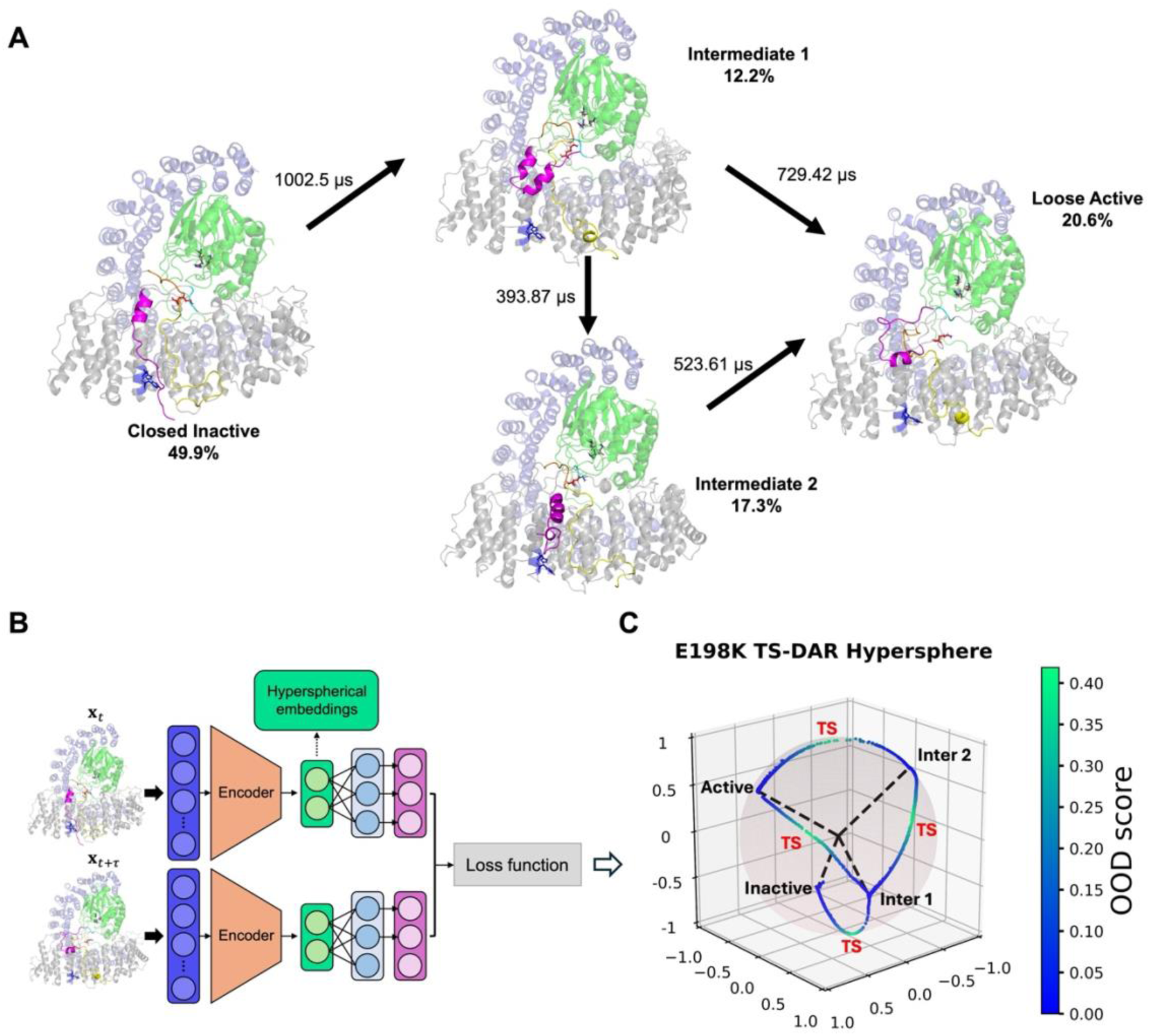
TS-DAR/MSM resolves four-state kinetics for E198K between closed inactive and loose active states. **(A)** Two major pathways connecting the four macrostates found during TS-DAR and MSM analysis with a closed inactive state, a loose active state and two intermediate states, labeled with the stationary populations for each state and MFPTs for each transition. The two major pathways had comparable flux: 56% including intermediate 2 and 44% bypassing intermediate 2. **(B)** Schematic of TS-DAR neural network architecture adapted from Liu et al. 2025.^50^ **(C)** Three-dimensional TS-DAR hypersphere embedding for E198K trained model. Each simulation frame is represented by a point on the sphere colored by the OOD score. The state centers and transition-state candidates are labeled.

An MSM was then constructed from these states using a lag time of 400 ns and validated through implied time scale (ITS) analysis and Chapman-Kolmogorov (CK) testing (Fig. S5).^30^ Transition path theory (TPT)^59,60^ identified two pathways with comparable flux connecting the closed inactive state, defined as the source, to the loose active state, defined as the sink. The dominant pathway proceeded from closed inactive to intermediate 1, then to intermediate 2, and finally to the loose active state, accounting for 56% of the total flux. The second pathway, representing 44% of the flux, bypassed intermediate 2 and transitioned directly from intermediate 1 to the loose active state. Mean first passage time (MFPT) analysis was used to estimate the timescales of each state transition (Table S2). For both pathways, the rate-determining step was the transition from the closed inactive state to intermediate 1, with an MFPT of 1,002.5 μs. These pathways allowed us to determine the conformational mechanism involved in the transition to the active state.

### Active state loosens N/C-arm interface rather than fully opening

The structure of the active state, in which both the active site and substrate-binding pocket are exposed, reveals a loosening of the N/C-arm interface that contrasts with an intuitive model of the activation mechanism. Based on the closed E198K structure, one might expect activation to occur through complete opening of the N/C-arm interface, fully exposing the catalytic residues. Instead, we found that this interface loosens while remaining associated with the holoenzyme core yet still providing access to the active site. During this process, the core remains the same between the inactive and active states, consistent with the loose and closed form cryo -EM structures.

Figure 3B-E shows bar plots for the key conformational features described in Figure 1, grouped by the states identified in Figure 2A. For the two active-site distances (“Active Site to N-arm” and “Active Site to C-arm”), the fraction of conformations with distances greater than 20 Å is shown for each state (Fig. 3B and C). For the “C-arm SLiM” and “N-arm Tail” distances, the average distance is reported (Fig. 3D and E). In the closed inactive state, the active site is occluded by the N/C-arm interface, as indicated by the low fractions of open conformations, and the substrate-binding pocket remains inaccessible due to the low “C-arm SLiM” distance, rendering the enzyme inactive. In intermediate 1, the N/C-arm interface becomes perturbed, with the N-arm exhibiting a higher fraction of open conformations while the C-arm remains largely closed. At the same time, the substrate-binding pocket becomes exposed, as indicated by the large average “C-arm SLiM” distance. Interestingly, in intermediate 2, the N- and C-arms appear to switch. The C-arm exhibits a greater fraction of open conformations than in intermediate 1, while the N-arm becomes more closed. In addition, the C-arm SLiM appears to partially backtrack, resulting in a decrease in the average “C-arm SLiM” distance relative to intermediate 1. In the loose active state, both the N- and C-arms display high fractions of open conformations, indicating an exposed active site, and the large “C-arm SLiM” distance indicates an exposed substrate-binding pocket. As a geometric check, we additionally docked the hexapeptide used for the *in vitro* phosphatase assay, reproduced in Figure 4B, into the active state structure using AutoDock Vina (see Methods Section K for details).^61,62^ We found that the exposed active site pocket of the active can accommodate the phosphorylated hexapeptide (Fig. S8). Together, these features are consistent with a catalytically competent state. The “N-arm Tail” distance remains relatively similar among the closed inactive, intermediate 1, and intermediate 2 states but becomes substantially more flexible in the active state, as reflected by the larger standard deviation (Fig. 3E).

**Figure 3.**
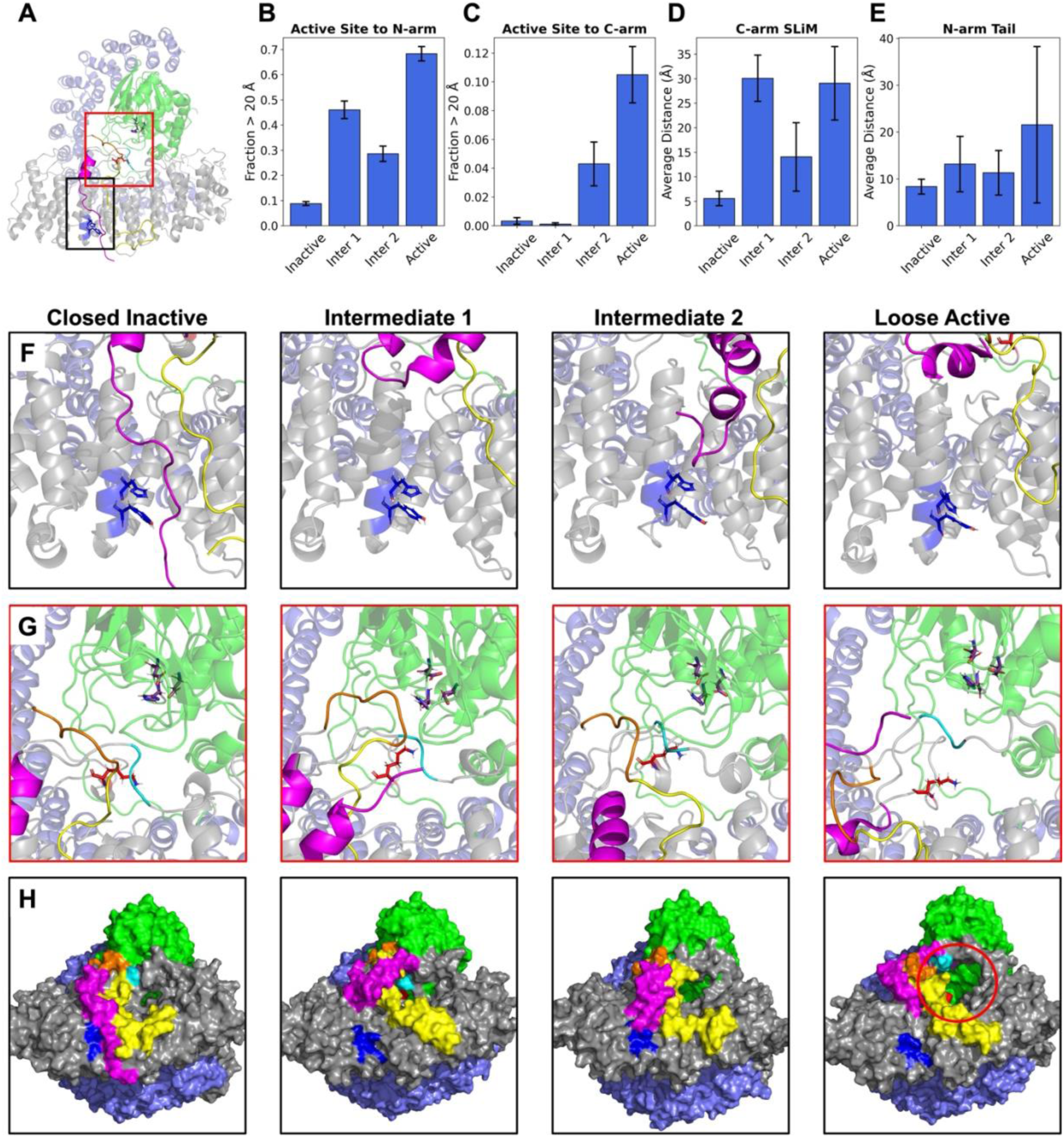
Four E198K macrostates resolved by TS-DAR show structural distinctions along the conformational pathway from closed inactive and loose active states. **(A)** Full view structure with zoomed in views highlighted. **(B)** Bar plot of fraction of “Active Site to N-arm” distances > 20 Å for each state in Figure 2A. **(C)** Same plot as (B) but for “Active Site to C-arm” distance. **(D)** Average “C-arm SLiM” distance for each state. **(E)** Same plot as (D) but for “N-arm Tail” distance. Distances are defined in Figure 1. Error bars for the fraction bar plots (C, B) are standard deviations from bootstrapping over the trajectories. Error bars for the average bar plots (D, E) are the standard deviation of the average distances. **(F)** Zoomed-in view of substrate-binding pocket and C-arm SLiM for each macrostate. **(G)** Zoomed-in view of the N/C-arm interface and active site for each macrostate. **(H)** Full-structure surface view showing substrate-binding pocket and path to active site. Residue colors correspond to those shown in Figure 1.

**Figure 4.**
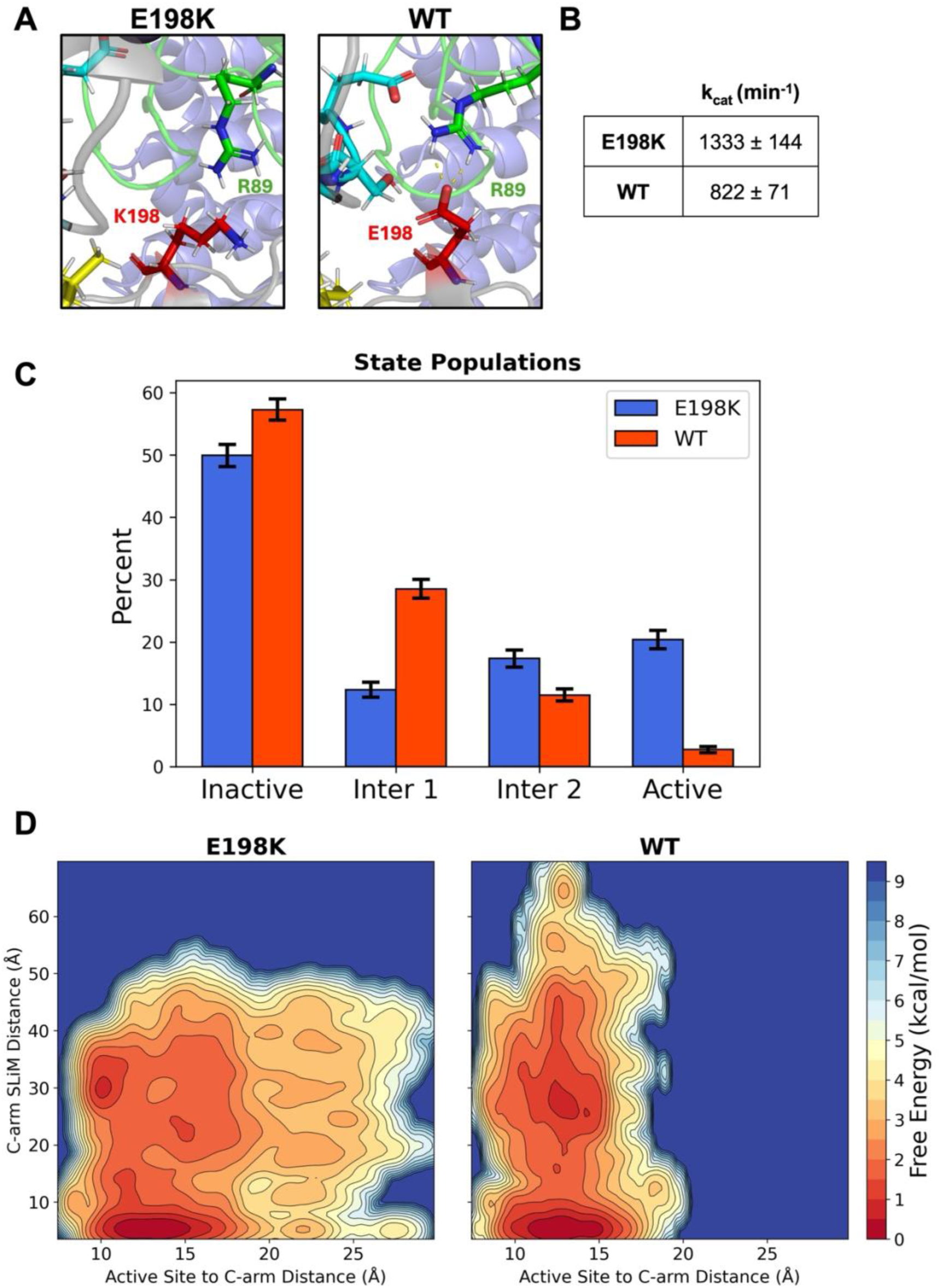
E198K has less occluded active site compared to WT resulting in hyperactivity for disease-associated variant. **(A)** View of mutation site residue 198 and catalytic subunit R89 interaction from closed inactive structure. E198K (left) has a broken salt bridge and WT (right) has a formed salt bridge. **(B)** Catalytic constants (k_cat_) for E198K and WT reproduced from Konovalov et al. 2023.^23^ **(C)** State populations for the macrostates shown in Figure 2A for E198K from TS-DAR model and WT from the closest E198K state center (see Methods Section H for details). **(D)** Two-dimensional FES for the “C-arm SLiM” distance (y-axis) and the “Active Site to C-arm” distance (x-axis) calculated from aggregated unbiased MD simulations.

We next examined the structural features of the TS-DAR hyperspherical embedding state centers (Fig. 2C), ordered along the dominant activation pathway. Figure 3F shows a zoomed-in view of the substrate-binding pocket (blue) for each state. In the closed inactive state, the C-arm SLiM (magenta) covers the binding pocket and blocks substrate binding. In contrast, the binding pocket is exposed in the remaining three states. Although the C-arm SLiM distance decreases from intermediate 1 to intermediate 2, the structural view indicates that the pocket remains largely accessible. Figure 3G shows the N/C-arm interface that obstructs access to the catalytic residues (purple). In the closed inactive state, both the N- and C-arms block the path to the active site. In intermediate 1, the N-arm shifts upward and adopts a kinked conformation while the C-arm remains in place. In intermediate 2, the C-arm instead shifts upward while the N-arm maintains coverage of the active site. In the active state, the N-arm moves furthest from the catalytic site, allowing the C-arm to shift upward and expose the active site. Full surface representations of each state are shown in Figure 3H. Both the active site and substrate-binding pocket are clearly occluded in the closed inactive state, whereas only the active site is occluded in intermediate 1 while the substrate-binding pocket is exposed. Although the catalytic site begins to open in intermediate 2, residual N/C-arm interactions continue to partially obstruct the route from the substrate-binding pocket to the catalytic center. Only in the loose active state are both regions fully accessible. Together, these structural analyses demonstrate that activation proceeds through progressive loosening of the N/C-arm interface rather than complete opening, exposing both the substrate-binding pocket and catalytic site while preserving the overall holoenzyme architecture.

Based on our MSM analysis, we suggest that PP2A-B56δ activation proceeds through two major pathways. In the first pathway, the enzyme begins in the closed inactive state with both the active site and substrate-binding pocket occluded. The N-arm covering the active site first loosens while the C-arm SLiM dissociates, exposing the substrate-binding pocket and producing intermediate 1. Next, partial backtracking of the N-arm and C-arm SLiM occurs, resulting in intermediate 2, where the substrate-binding pocket remains exposed but active-site accessibility is reduced by the N-arm. Instead, in intermediate 2, the C-arm covering the active site starts to loosen, followed by coordinated loosening of the N-arm, while the C-arm SLiM moves further away from the binding pocket. These collective motions expose both the active site and substrate-binding pocket, producing the fully activated state. In the second pathway, the enzyme bypasses intermediate 2 and transitions directly from intermediate 1 to the active state. In this pathway, activation proceeds through sequential loosening of the N-arm and displacement of the C-arm SLiM, followed by movement of the C-arm away from the active site, resulting in conversion from the closed inactive state to the loose active state. Although two activation pathways are observed, both converge on the same mechanistic principle: progressive loosening of the dual autoinhibitory N/C-arm interface sequentially exposes the substrate-binding pocket and catalytic site while preserving the overall holoenzyme architecture.

### Loose active state is more populated in E198K compared to WT

To understand the differences between E198K and WT, we assigned each WT conformation to the closest E198K state center and classified it according to the corresponding E198K macrostate (see Methods Section H for details). A comparison of the resulting state populations is shown in Figure 4C. The largest differences between the two variants occur in the intermediate 1 and active states with smaller differences for inactive and intermediate 2. In particular, the active state is more populated in E198K than in WT. This increased active-state population is consistent with previous experimental measurements showing that E198K exhibits a higher k_cat_ than WT (reproduced in Fig. 4B).^23^ To further compare the conformational landscapes of the two variants, we constructed a two-dimensional free energy surface (FES) from the unbiased MD simulations using the “Active Site to C-arm” distance (x-axis) and the “C-arm SLiM distance” (y-axis) as collective variables (Fig. 4D). The FES reveals that E198K samples a larger population of conformations with a more exposed active site compared to WT.

Taken together, the increased population of exposed active-site conformations and the higher occupancy of the active macrostate indicate that E198K shifts the conformational equilibrium toward more active states. These observations are consistent with experimental measurements and support the conclusion that E198K is a hyperactive variant relative to WT. In contrast, WT exhibits a larger population of intermediate 1, a state characterized by an exposed substrate-binding pocket but limited accessibility to the active site. This observation is consistent with the FES, which shows that WT samples larger “C-arm SLiM” distances than E198K. However, as shown in Figure 3H, exposure of the substrate-binding pocket alone is insufficient for activation; both the substrate-binding pocket and active site must be accessible for productive catalysis. The shift toward active conformations observed in E198K may underlie the molecular basis of its pathogenic role in intellectual disability.

Inspection of the mutation site reveals that E198K disrupts a salt bridge between residue 198 of the regulatory subunit and R89 of the catalytic subunit that is present in the WT enzyme (Fig. 4A). This R89-E198 salt bridge has been observed in previous literature and a quantum mechanics study found that R89 binding of substrate phosphate is critical for catalytic efficiency.^24,63^ Salter and colleagues hypothesized that this salt bridge could help regulate the catalytic activity of catalytic subunit. To further understand how disruption of this salt bridge promotes activation of the holoenzyme, we next examined the candidate transition-state ensembles connecting the macrostates.

### E198K affects metastable state stability rather than disrupting the transition states

TS-DAR can simultaneously identify candidate transition-state ensembles between all metastable macrostates.^50^ These candidates correspond to conformations with high OOD scores that lie between metastable states on the hyperspherical embedding (Fig. 2C). However, candidate structures must be validated through unbiased simulations to determine whether they represent true transition states. A true transition state is expected to exhibit approximately equal commitment probabilities to the neighboring macrostates. To identify transition-state candidates for each macrostate pair, we selected the 100 conformations with the highest OOD scores and clustered them into five groups (see Methods Section J for details). From each cluster, the conformation with the highest OOD score was chosen as a representative candidate and used to initiate 20 independent 50 ns unbiased simulations. Each trajectory was subsequently analyzed using the trained E198K TS-DAR model to assign macrostate labels. For each transition, the candidate structure exhibiting the most even distribution of state assignments, based on the final 5 ns of the trajectories, was selected as the putative transition state (Fig. S7). These structures were then reverted to WT and subjected to the same procedure.

A comparison of the resulting state populations for the WT and E198K transition-state structures is shown in Figure 5A. Overall, only minor differences were observed between the two variants, suggesting that the E198K mutation has little effect on the transition state connecting the metastable macrostates. The little to no effect on the transition state implies that E198K does not change any activation free energy barrier along the activation pathway. When combined with the state population analysis presented in Figure 4C, these results suggest that the primary effect of the E198K mutation is not to alter the transition states themselves but rather to shift the relative stability of the metastable states. Specifically, the WT enzyme appears to preferentially stabilize the closed inactive state and intermediate 1, thereby reducing the likelihood of transitioning into the active state compared to E198K. Together, these results suggest that E198K promotes PP2A-B56δ activation primarily by shifting the conformational equilibrium toward active states rather than by lowering the activation free energy barriers between them.

**Figure 5.**
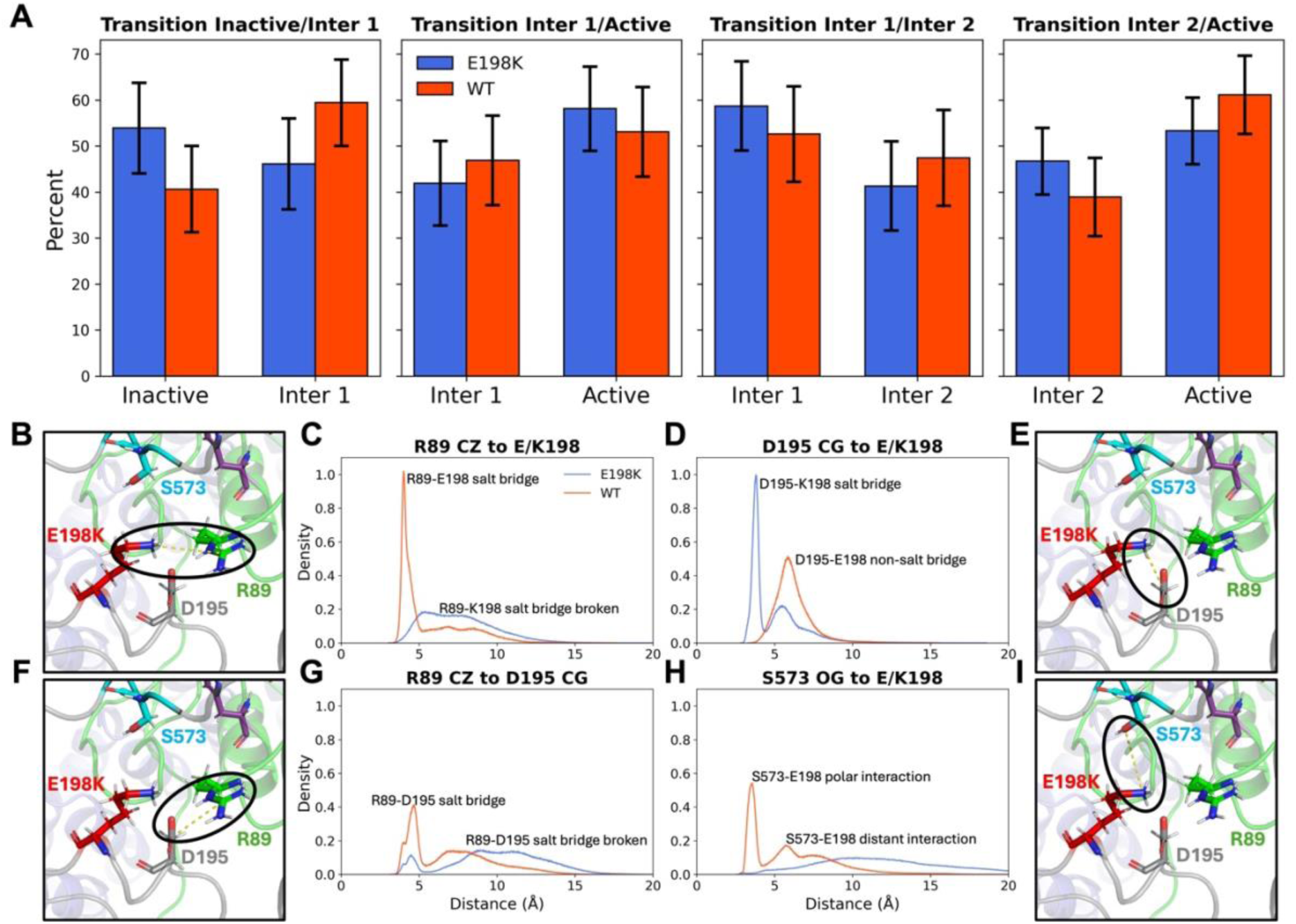
E198K does not affect transition states but instead weakens interactions with C- arm allowing exposure of active site. **(A)** Bar plots comparing E198K and WT state populations from the 20 x 50 ns transition-state structure simulations for each pair of macrostates. Error bars are the standard deviation from bootstrapping the 20 trajectories. **(B-I)** Raw data histograms of specific atomic distances comparing the E198K and WT full dataset of simulations. Each histogram is accompanied by an E198K structural image of the internal loop and C-arm interface. **(B, C)** Catalytic site residue R89 atom CZ to E/K198 atom CD/NZ distance. **(D, E)** Regulatory subunit internal loop residue D195 atom CG to E/K198 atom CD/NZ distance. **(F, G)** R89 atom CZ to D195 atom CG distance. **(H, I)** Regulatory subunit C-arm residue S573 to E/K198 atom CD/NZ distance.

### E198K weakens interaction with C-arm covering active site resulting in overactivity

To understand the structural origin of this shift in conformational equilibrium, we next examined the interaction network surrounding the E198K mutation. As shown in Figure 4A, the E198K mutation disrupts a salt bridge between residue 198 of the regulatory subunit and R89 of the catalytic subunit that is present in the WT enzyme. This salt bridge is part of a broader network of interactions involving the internal loop, which contains the mutation site, the catalytic subunit, and the C-arm that covers the active site. Disruption of this interaction network weakens contacts between the internal loop and the C-arm, allowing the C-arm to move more freely away from the active site. Increased mobility of the C-arm promotes exposure of the catalytic site and contributes to the higher active-state population observed for E198K relative to WT.

Figures 5B-I show histograms of key residue-residue distances within this interaction network with structural images of the corresponding distances. The histograms represent the raw populations observed across the aggregate unbiased simulations for WT and E198K. The first structural view and histogram (Fig. 5B and C) show the distance between the CZ atom of R89 and the distal heavy atom of residue 198, directly representing the R89-E/K198 interaction. In WT, the distribution exhibits a sharp peak near 4 Å, consistent with a stable salt bridge.^64^ In contrast, the E198K distribution is much broader and lacks a pronounced peak at this distance, indicating disruption of the salt bridge. Following the loss of the R89-E198 interaction, K198 in the mutant can instead form a salt bridge with D195, another residue within the internal loop (Fig 5E). Consistent with this interaction, Figure 5D shows a peak near 4 Å for E198K. In WT, the corresponding distribution is shifted to larger distances and resembles a broad Gaussian distribution, which is not indicative of a salt bridge but rather simple proximity of these residues. D195 can also interact with R89, although this interaction is substantially less populated than the primary salt bridges and is observed less frequently in E198K than in WT (Fig. 5F and G).

The key interaction linking the mutation site to active-site accessibility is between S573 of the C-arm and residue 198 (Fig 5I). As shown in Figure 5H, this distance exhibits a multimodal distribution with a prominent peak near 4 Å for WT, corresponding to H-bonding, while E198K exhibits a much broader distribution. These results indicate that the S573-E198 interaction is substantially more stable in WT than the corresponding S573-K198 interaction in E198K. Based on this network of interactions, we propose that formation of the K198-D195 salt bridge competes with and weakens interactions between residue 198 and S573. As a result, the C-arm is less tightly coupled to the internal loop and can more readily move away from the active site, increasing active-site exposure and favoring the active state. In WT, the stronger interaction between E198 and S573 helps maintain C-arm coverage of the catalytic site, thereby stabilizing inactive conformations. The maintenance of this C-arm coverage in WT is consistent with its increased population of intermediate 1, which has a low fraction of open C-arm (Fig. 3C) and decrease of intermediate 2, which has a higher fraction, based on the populations shown in Figure 4C.

Interestingly, S573 is a known phosphorylation site in PP2A-B56δ according to PhosphoSitePlus,^65^ and phosphorylation at this site has been previously shown to increase enzymatic activity.^17,23^ In mouse models, phosphorylation by protein kinase A of residue S566, corresponding to human S573, was found to be associated with enzyme activation.^66^ The mechanism proposed here provides a structural explanation for this observation. Introduction of a negatively charged phosphate group at S573 would create unfavorable electrostatic interactions with E198 in WT, weakening the S573-E198 contact and promoting displacement of the C-arm from the active site. Thus, both phosphorylation of S573 and the E198K mutation may activate PP2A-B56δ through a common mechanism involving disruption of the interaction network that stabilizes C-arm coverage of the active site. Therefore, the disease-associated E198K mutation appears to mimic the activating effect of S573 phosphorylation by destabilizing interactions that maintain C-arm coverage of the active site.

## Overall Discussion

Our cryo-EM analysis identified two conformations of the PP2A-B56δ disease-associated mutant E198K with identical holoenzyme cores. One structure had disordered N/C-arms, referred to as the active loose-form. The other structure had less flexibility of this region allowing for the N/C-arms to be resolved, called the inactive closed-form. In the closed-form, the active site and substrate binding pockets were blocked by the N/C-arms resulting in an inactive structure. Conversely, in the loose form, the N/C-arm could not be resolved and these pockets were exposed. Particle analysis showed that the loose form was more prevalent compared to the closed form for E198K. Additionally, the loose form had a higher population compared to a previously resolved set of structures of PP2A-B56δ E197K variant indicating higher activity, consistent with experimental results.^17^

We characterized the activation mechanism of the PP2A-B56δ holoenzyme and found that activation proceeds through sequential loosening of the uniquely long N- and C-arms that mediate the dual autoinhibitory architecture of this regulatory subunit, while preserving the overall holoenzyme architecture. To our knowledge, the activation mechanism of a dual-autoinhibited phosphatase of this nature has not been previously described. The activation of the holoenzyme occurred by a coordinated loosening of the N/C-arm interface rather than a complete opening. A complete unfolding of the interface would disrupt the overall architecture of the regulatory subunit creating more disorder in the N- and C-arms resulting in a large entropic barrier for the reformation of the interface. We show that loosening rather than full opening prevents this entropic barrier from forming allowing for a reversible conformational change that regulates the activation of the enzyme. This regulation is key to maintaining homeostasis of cellular phosphorylation levels.

The proposed activation mechanism is unique to B56δ because of this isoform’s significantly longer N- and C-arms compared to others in its family.^17^ The loosening mechanism identified here suggests that activation can occur without complete displacement of the N/C-arm interface, allowing the holoenzyme to retain its overall structure while exposing the substrate-binding pocket and catalytic site. This structural arrangement may have important implications for substrate recognition. Because access to the catalytic site remains partially constrained in the loosened state, substrates may need to follow a less direct path from the SLiM-binding pocket to the active site than would be allowed in a fully open conformation. This indirect path would allow the B56δ isoform to accommodate a longer spacer length between the SLiM motif and the phosphorylation site. Previous bioinformatics studies have found that the B56 regulatory subunit family has longer spacer lengths compared to the B55 family which could be due to the contribution of the B56δ isoform.^67,68^ Additionally, these substrates could interact with portions of the N/C-arms which could change their disorder structure while binding a substrate. Future studies examining substrate recognition across PP2A regulatory subunits will be important for determining whether the unique architecture and activation mechanism of B56δ contribute to distinct substrate preferences and specialized signaling functions.

The cryo-EM particle fractions should not be directly compared with the equilibrium populations predicted by the MSM. Although cryo-EM samples the solution conformational ensemble, the relative particle fractions obtained after sample preparation and image-processing procedures are not expected to quantitatively preserve the equilibrium populations of the underlying conformational states.^69^ Consequently, the cryo-EM particle fractions should be interpreted qualitatively rather than as Boltzmann populations.

The comparison between WT and E198K found that active state was more populated and the catalytic site was more exposed for E198K compared to WT, suggesting the disease-associated mutation preferentially adopts active conformations and is thus more catalytically active. This finding is consistent with previous biochemical studies reporting increased catalytic activity for E198K.^15,17,23^ This variant’s hyperactivity would cause increased levels of dephosphorylation thus prematurely deactivating enzymes in critical signal transduction pathways like CREB (cAMP Response Element-Binding protein) signaling.^1^ Disruption of this pathway can cause issues for neurological function and memory formation.^70–72^ Interestingly, transition state analysis showed that E198K did not significantly affect the transition state between metastable states, indicating that the mutant does not lower the activation free energy barrier along the conformational activation pathway. Instead, E198K appears to promote activation by destabilizing inactive conformations and shifting the conformational equilibrium toward pre-existing active states. These findings support a conformational selection mechanism in which the mutation remodels the free-energy landscape without fundamentally changing the activation pathway.

Our interaction network analysis uncovered atomistic details explaining this shift in equilibrium conformational populations and identified key residues involved in the stabilization of the inactive state. Residue R89 of the catalytic subunit has been previously shown to coordinate with the substrate phosphate group resulting in an important interaction for efficient catalysis.^63^ We found that the E198-R89 salt bridge in the WT system was broken in E198K by the introduction of the positively charged LYS residue. This salt bridge was previously hypothesized to help regulate the enzyme by preventing R89 from interacting with substrates.^24,63^ Consequently, breaking this salt bridge in E198K allows this residue to interact with the phosphorylated substrate resulting in dephosphorylation. Our results further reveal that the effects of E198K extend beyond this single interaction and instead involve a broader network of contacts linking the internal loop to the C-arm that occludes the catalytic site. In WT, residue E198 forms a stabilizing interaction with the regulatory-subunit phosphorylation site S573, helping maintain coupling between the internal loop and the C-arm, thereby promoting catalytic-site occlusion. In E198K, formation of an additional K198-D195 salt bridge competes with this interaction and weakens coupling between residue 198 and S573. The resulting increase in C-arm mobility favors exposure of the catalytic site and shifts the conformational equilibrium toward active states. The phosphorylation of S573 would be expected to disrupt the E198-S573 interaction through electrostatic repulsion, providing a potential mechanistic explanation for the activating effect of S573 phosphorylation. We therefore propose that disruption of the internal loop-C-arm interaction network represents a canonical mechanism for activation of PP2A-B56δ.

Several important questions remain for future investigation. One promising direction is to examine substrate binding within the transition-state ensembles and metastable states identified here to further understand how phosphorylated substrates interact with the dynamic autoinhibitory architecture of PP2A-B56δ. For example, substrate binding may preferentially stabilize certain transition-states, such as the transition between intermediate 1 and the active state, thereby shifting the conformational equilibrium toward activation. Structural modeling and substrate-docking studies could provide valuable insight into how substrate recognition is coupled to the activation mechanism proposed in this work.

## Conclusion

In this study, we combined cryo-EM, enhanced-sampling MD simulations, TS-DAR analysis, and MSMs to elucidate the activation mechanism of PP2A-B56δ. Our cryo-EM analysis identified an inactive closed form and an active loose form of E198K PP2A-B56δ, revealing that activation is governed by conformational rearrangements of the N/C-arms but leaving the underlying dynamic mechanism unresolved. Large-scale MD simulations combined with TS-DAR and MSMs revealed four metastable states connected by two dominant activation pathways. We found that activation does not occur through complete opening of the N/C-arm interface, as might be intuitively expected, but through progressive loosening of the N- and C-arms that exposes both the substrate-binding pocket and catalytic site while maintaining the overall holoenzyme structure. Comparison of the E198K mutant with WT showed that the disease-associated mutation promotes activation primarily by shifting the equilibrium populations of the metastable states while leaving the transition-state ensemble largely unchanged. Our simulations further identified an interaction network involving the internal loop, R89, D195, E198, and the C-arm that regulates C- arm mobility and provides an atomistic explanation for the hyperactivity of the E198K mutation. Altogether, our findings establish progressive loosening of the dual autoinhibitory interface as the molecular basis for PP2A-B56δ activation and demonstrate how integrating cryo-EM with MD simulations and kinetic modeling can uncover conformational mechanisms and allosteric regulation in highly dynamic protein complexes.

## Methods

### A. Protein expression and purification

All constructs were generated using standard polymerase chain reaction (PCR)-based molecular cloning. Human PP2A scaffold A subunit (Aα) carrying an N-terminal His6 tag, PP2A catalytic subunit (Cα) with an N-terminal His8 tag, and GST-tagged B56δ were expressed in insect cells using a Bac-to-Bac baculovirus expression system. The E198K mutation of B56δ subunit was generated by site-directed mutagenesis.

For holoenzyme expression, Hi-5 cells were grown to a density of 1.5 × 10^6^ cells/mL and co-infected with baculoviruses encoding His-tagged scaffold A subunit, His-tagged PP2Ac, and GST-tagged B56δ. After 48 h of expression at 27 °C, cells were harvested and lysed by Dounce homogenization in buffer containing 25 mM Tris-HCl (pH 8.0), 150 mM NaCl, 50 μM MnCl_2_, 2 mM DTT, and protease inhibitors (10 μM leupeptin, 0.5 μM aprotinin, and 1 mM PMSF). Cell debris and insoluble material were removed by centrifugation, and the clarified lysate was applied to glutathione Sepharose 4B resin (Cytiva) three times. The resin was subsequently washed twice with 5 column volumes of lysis buffer. To remove the GST tag, TEV protease was added directly to the resin and incubated for 16 hours at 4 °C. The untagged holoenzyme was collected in the flow-through and further purified by anion-exchange chromatography using a Source 15Q column (Cytiva), followed by size-exclusion chromatography on a Superdex 200 column (Cytiva) (Fig S1C).

### B. Cryo-EM sample preparation and data acquisition

Cryo-EM grids of the E198K B56δ holoenzyme were prepared at the New York Structural Biology Center using a prototype Chameleon system (SPT Labtech) based on Spotiton technology.^73,74^ Briefly, 50 pL of holoenzyme sample at 1.6 mg mL^-1^ was dispensed onto homemade self-wicking nanowire grids^75^ and immediately plunge-frozen in liquid ethane. Grid preparation was carried out at room temperature under moderate humidity conditions without humidity control.

Cryo-EM data for the E198K holoenzyme were collected on a Titan Krios electron microscope (Thermo Fisher Scientific) operated at 300 kV and equipped with a Gatan K2 Summit direct electron detector. A total of 2,519 movies were acquired automatically using Leginon^76^ over a defocus range of −1.2 to −2.0 μm at a nominal magnification of 105,000×, corresponding to a calibrated pixel size of 1.096 Å/pixel. Each movie was dose-fractionated into 50 frames, with a total accumulated electron exposure of 66.84 e^-^/Å^2^ (Table S1).

### C. Cryo-EM data processing

Image processing was carried out in cryoSPARC v4.0^77^ and the workflow is summarized in Figure S1 and Table S1. Movie stacks were first corrected for beam-induced motion using Patch Motion Correction, and contrast transfer function (CTF) parameters were estimated with Patch CTF Estimation. Micrographs with estimated CTF resolutions worse than 4 Å were excluded from further analysis. Initial particle picking was performed on 30 high-quality micrographs using Blob Picker. The resulting particles were subjected to 2D classification, and well-defined classes representing multiple particle orientations were used as templates for automated particle picking across the full dataset. Extracted particles were boxed at 280 Å and subjected to three rounds of 2D classification to remove false positives and damaged particles. The curated particle set was then used for *ab initio* reconstruction, followed by heterogeneous and homogeneous refinement. To resolve distinct structural states of the E198K PP2A-B56δ holoenzyme, 3DVA was performed.^51^ Particles assigned to conformationally homogeneous classes were further refined by homogeneous refinement, followed by both global and local CTF refinement. For the closed-state reconstruction, local refinement was additionally performed using a mask focused on the B56δ subunit to improve local map quality. These procedures yielded reconstructions at 2.93 Å resolution for the loose conformation and 3.20 Å resolution for the closed conformation of the E198K PP2A-B56δ holoenzyme (Fig S1A and B).

### D. Model building and refinement

Initial models of the PP2A-B56δ holoenzyme was generated using the structures of the loose-form (PDB ID: 8U1X) and closed-form (PDB ID: 8U89) E197K PP2A-B56δ complexes as starting templates.^17^ The models were manually docked into the cryo-EM density using UCSF Chimera,^78^ and model adjustment and rebuilding were carried out in Coot.^79^ Final refinements were performed in PHENIX using the phenix.real_space_refine module with secondary structure and geometry restraints applied throughout refinement.^80^ The model refinement statistics are summarized in Table S1.

### E. Initial Structure and Equilibration Simulation Setup

Our novel cryo-EM structure of the closed-form E198K variant was used to initialize the all-atom MD simulations. Residues 49-58, 513-564, and 602 of the regulatory subunit B56δ were filled in using MODELLER^52^ integrated in ChimeraX.^81–83^ The resulting protein consisted of residues 8-589 for the scaffold subunit A (UnitProt: P30153),^84^ 49-602 for the regulatory subunit B56δ (UnitProt: Q14738-1),^85^ and 2-309 for the catalytic subunit (UnitProt: P67775),^86^ plus two manganese ions from the cryo-EM structure. Using PyMOL,^87^ residue L309 of the catalytic subunit was methylated because this methylation is present during the assembly of the heterotrimeric holoenzyme in cells.^88,89^

The structure was then solvated in a dodecahedron box with 71,572 waters, using the TIP3P water model.^90,91^ Sodium and chloride ions were added to neutralize the system and achieve a physiological salt concentration of 0.15 M. The total number of atoms used for the simulation box was 238,467 atoms. The MD simulations were run using GROMACS 2022.5^92,93^ with the AMBER14SB force field.^94^ The Lennard-Jones force fields for the Mn ions were adopted from Bradbrook et al. 1998.^95^ The neighbor list, Lennard-Jones interactions, and Coulomb interactions were all cut off at 12 Å and the long-range interactions were computed with particle-mesh Ewald.^96^ The system was energy minimized using the steepest descent algorithm, then equilibrated for 1 ns in the NVT ensemble, followed by 1 ns in the NPT ensemble with position restraints on the protein heavy atoms using a force constant of 1000 kJ/mol. Following the equilibration, a 20-ns unrestrained NVT simulation was performed to further relax the system before enhanced sampling. The simulations were performed at 300 K maintained with the V-rescale thermostat using a 0.1-ps time constant^97^ and the pressure was maintained at 1 bar using the Parrinello-Rahman barostat with a coupling constant of 2.0 ps and isotropic coupling.^98^ The leapfrog integrator was used for equilibration and production with a 2-fs timestep using LINCS to constraint the hydrogen atom bonds.^99^ The same process was used for the WT system which we reverted from the E198K initial structure using PyMOL.^87^ The total number of atoms for the WT system was 238,450 atoms.

### F. Enhanced Sampling Simulation Setup

After the equilibration setup, we employed replica exchange with solute tempering (REST2) simulations to enhance the conformation sampling of the protein system.^53–55^ We used residues 49-78 (N-arm) and 580-602 (C-arm) of the regulatory subunit as the Hamiltonian-scaled region (“hot region”), comprising of 816 atoms across 53 residues. A total of 16 replicas were run for 600 ns each, with Hamiltonian scaling factors ranging from 1.0 to 0.5 (effective temperature of 300 to 600 K). Exchanges between neighboring replicas were attempted every 1 ps and accepted according to the Metropolis criterion. The simulations were performed using GROMACS 2022.5^92,93^ patched with PLUMED 2.9.^100^

### G. Large-scale MD Simulation Setup and Analysis

The REST2 trajectories were projected onto the “C-arm SLiM” and “Active Site to C-arm” distances, described in Figure 1. We then clustered the space using K-means^56^ and chose the conformations closest to the cluster centers as starting conformations for Folding@Home seeds.^57^ After excluding structures in which the Mn ions had dissociated from the catalytic site, 176 initial structures were retained for E198K and 176 for WT. The Folding@Home simulations were performed using OpenMM 8.1.1 in the NVT ensemble with the Langevin middle integrator^101^ with a 2-fs timestep and a friction coefficient of 1.0 ps^-1^.^102^ After excluding trajectories shorter than 100 ns, a total of 859 trajectories were collected for E198K and 874 for WT, corresponding to aggregate simulation times of 1.04 ms and 1.17 ms, respectively (Fig. S2). From these simulations, we measured the pairwise distances between the Cα’s of residues 61-90, 180-312, 560-602 of the regulatory subunit B56δ and residues 55-60, 84-92, 239-246, 260-270 for a total of 240 residues yielding 28,680 pairwise distances. These residues were chosen based on prior structural knowledge and their involvement in holoenzyme activation. All distance measurements were conducted using MDAnalysis 2.7.0.^103,104^

### H. TS-DAR Model Training

From the 28,680 pairwise Cα–Cα distances, we used Spectral-oASIS^58^ to select 1,000 features to train a four-macrostate TS-DAR model. Unless otherwise noted, all training parameters were identical to those reported in the original TS-DAR methodology publication.^50,105^ The training lag time was 10 ns, the batch size was 1,000, and the neural network architecture consisted of layers with dimensions [1000, 550, 250, 100, 50, 25, 3]. A schematic of the architecture is shown in Figure 2B, adapted from Liu et al. 2025.^50^ The dimensionality of the latent hyperspherical embedding was set to three for visualization and per recommendation of the original method when training a four-state model.^50,105^ Training consisted of 10 independent trials using different random initialization seeds. Each trial was trained for 10 epochs using both dispersion and VAMP-2 (variational approach for Markov processes)^106^ loss functions for all epochs. The losses were consistent over the independent training runs (Fig. S3). The final TS-DAR model was chosen based on the hypersphere with the most compact metastable states that had no isolated states (Fig. 2C). The representative structures shown in Figure 2A are the structures closest to the TS-DAR model state centers in the hyperspherical embedding space. For the WT system, state assignments were determined by identifying the nearest E198K state-center structure in the 1,000-feature space used to train the TS-DAR model.

### I. MSM Construction, Validation, and Analysis

From the final TS-DAR model, we used the state assignments to build an MSM to characterize the long-timescale dynamics of the conformational changes involved in activating the mutant holoenzyme. We built a four-state MSM using the transpose reversibility method to estimate the transition probability matrix (TPM). From the ITS analysis, the slowest timescales reach stable behavior with longer lag times (Fig. S5A). We selected a lag time of 400 ns for the final MSM and validated using the CK test (Fig. S5B).^30^ From this validated MSM, we calculated the MFPTs and stationary populations for the E198K system (Fig. 2A). All MFPTs are reported in Table S2. Using TPT analysis we determined the pathways connecting the inactive (source) and active (sink) states. This analysis found two pathways of comparable flux: inactive, inter 1, inter 2, active (56%) and inactive, inter 1, active (44%). All MFPT and TPT calculations were performed using MSMBuilder 2022.^107–110^

### J. Transition-state Validation

High OOD-score conformations identified by TS-DAR represent candidate transition-state ensembles; however, a high OOD score alone does not guarantee that a structure exhibits equal commitment probabilities to the adjacent metastable states.^50,105^ To validate these candidate transition states, we selected the 100 highest OOD-score conformations located between each pair of neighboring metastable states on the hyperspherical embedding (Fig. 2C). The corresponding 1,000-feature representations were projected onto the first four principal components calculated from the full 1.04 ms E198K trajectory dataset using principal component analysis (PCA).^111^ These conformations were then clustered into five groups using K-means clustering,^56^ and the highest OOD-score structure from each cluster was selected as a representative candidate.

From those chosen structures, we ran 20 x 50 ns unbiased MD simulations, then extracted the last 5 ns of each trajectory and used the trained TS-DAR model to assign metastable states. The ratios between the adjacent metastable states were calculated from each structure and the structures with the ratio closest to one were chosen as the validated transition-state structures for each pair of metastable state transitions (Fig. S7). Based on this criterion, cluster 2 was selected for the Intermediate 1/Inactive transition, cluster 3 for the Intermediate 1/Active transition, cluster 2 for the Intermediate 1/Intermediate 2 transition, and cluster 4 for the Intermediate 2/Active transition.

To compare transition-state behavior between variants, the validated E198K transition-state structures were reverted to WT using PyMOL. For each WT transition-state structure, 20 x 50 ns unbiased simulations were performed. Metastable-state assignments were determined using the same TS-DAR-based procedure described above, and the resulting state populations were compared with those obtained for E198K (Fig. 5A). Equilibration of the transition-state simulations followed the protocol described in Methods Section E. Production simulations were carried out in the NVT ensemble, with independent initial velocities assigned to each trajectory.

### K. Substrate Docking using AutoDock Vina

We used AutoDock Vina version 1.1.2 to dock the phosphorylated hexapeptide KRpTIRR into the loose active state of the E198K variant shown in Figure 2A. The state center structure identified by the TS-DAR hypersphere (Fig. 2C) was used for docking the substrate. To ensure the substrate would dock into the active site, we used box dimensions that encompassed residues 560-574 of the regulatory subunit B56δ, the active site residues D57, D59, and H85 of the catalytic subunit, and the Mn ions. We chose the hexapeptide to match the substrate used in the *in vitro* phosphatase assay that determined the catalytic constants for E198K and WT from Konovalov et al. 2023,^23^ reproduced in Figure 4B. The best fit docked structure had an affinity of -9.1 kcal/mol, shown in Figure S8.

## Author Contributions

Y.X. and X.H. designed the research; M.S.O. performed MD simulations; M.S.O. and Y.L discussed and analyzed the computational data; C.G.W performed the experiments; M.S.O., Y.L., C.G.W, Y.X., and X.H. wrote the paper.

## Competing Interest Statement

The authors declare no competing interests.

## Acknowledgements

We acknowledge the support from the Hirschfelder Professorship Fund and Vilas Associate Award from University of Wisconsin-Madison to X.H, the Research Forward Fund from the University of Wisconsin-Madison Office of the Vice Chancellor for Research with funding from the Wisconsin Alumni Research Foundation to X.H., M.O. acknowledges a fellowship from the National Institutes of Health (NIGMS, T32 GM130550). Y.X. and X.H. acknowledge the funding from NIH/NIGMS under award number R01GM14581.

## Data Availability

The MD simulation feature trajectories are available from Zenodo (10.5281/zenodo.21285351). The Python analysis code can be accessed on our GitHub repository (xuhuihuang/pp2a-loosening.git).

## Notes

### Competing Interest Statement

The authors have declared no competing interest.

