## Supplemental Material for "Progressive Loosening of a Dual Autoinhibitory Interface Activates PP2A-B56δ"

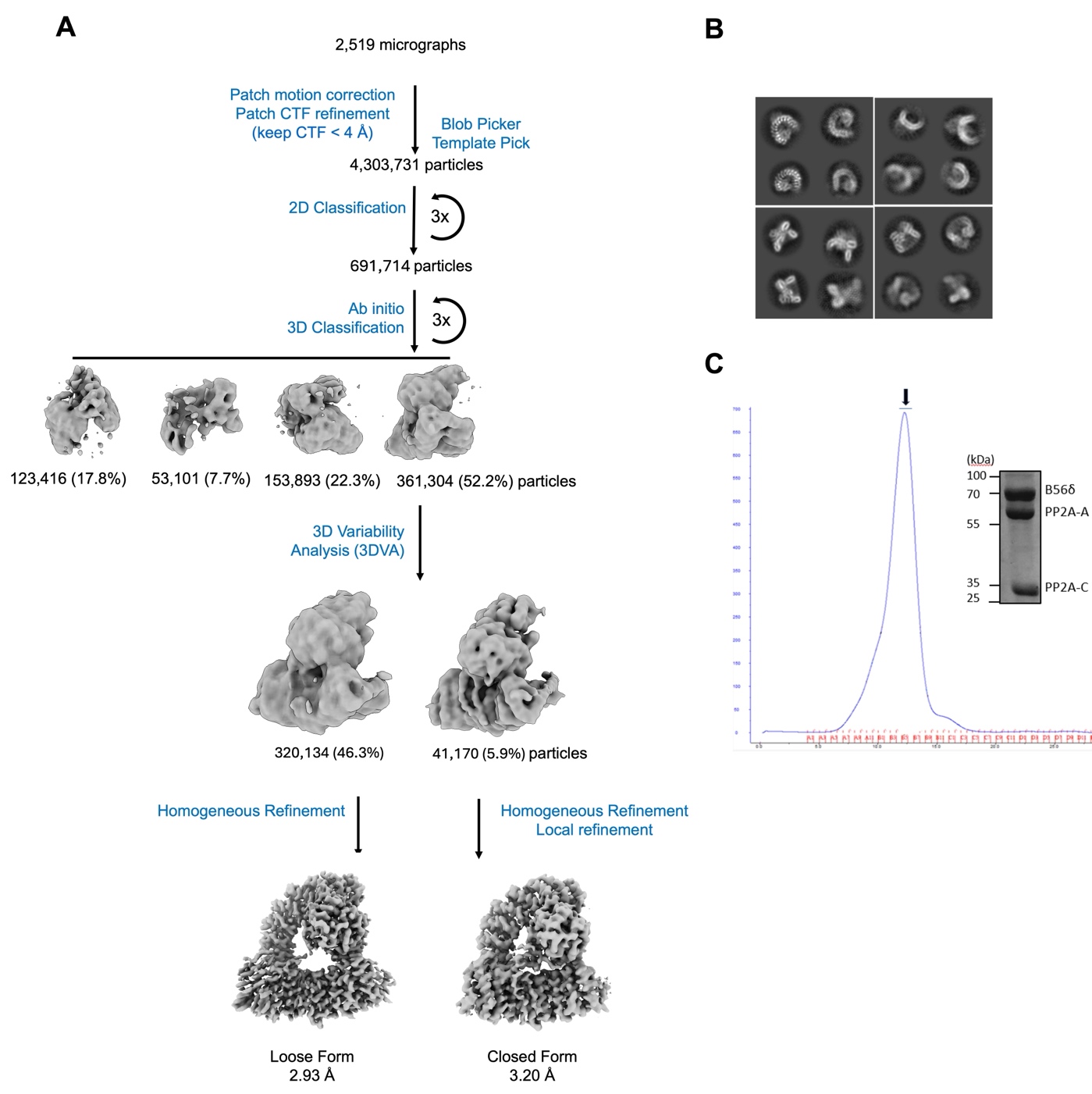
**Figure S1. (A)** Data analysis pipeline for E198K PP2A-B56δ cryo-EM structures showing two distinct forms, loose and closed. The closed form structure was used for MD simulations due the structural stability of the N/C-arms allowing for better resolution of this region compared to the loose form. **(B)** Representative 2D class averages of the E198K PP2A-B56δ holoenzyme. Particle percentages are based on total particles from 2D classification (691,714 particles). **(C)** Size-exclusion chromatography (SEC) chromatogram and sodium dodecyl dulfate-polyacrylamide gel electrophoresis (SDS-PAGE) analysis of the E198K PP2A-B56δ holoenzyme.

**Table S1. Cryo-EM data collection, refinement and validation statistics**

| PP2A–B56δ holoenzyme | E198K Closed Form | E198K Loose Form |
| --- | --- | --- |
| **Data collection and processing** | |  |
| Magnification (kx) | 105 | 105 |
| Voltage (kV) | 300 | 300 |
| Defocus (μm) | -1.2 to -2.0 | -1.2 to -2.0 |
| Pixel size (Å) | 1.096 | 1.096 |
| Total dose (e^-^ / Å^2^) | 66.84 | 66.84 |
| Number of micrographs | 2519 | 2519 |
| Number of frames | 50 | 50 |
| Number of initial particles picked | 4,303,731 | 4,303,731 |
| Number of final particles refined | 41,170 | 320,134 |
| Map resolution (Å) | 3.20 | 2.93 |
| FSC threshold (Å) | 0.143 | 0.143 |
| **Refinement** | |  |
| Non-hydrogen atoms | 10647 | 9944 |
| Protein residues | 1344 | 1262 |
| Ligand | MN:2 | MN:2 |
| RMSD |  |  |
| bond length (Å^2^) | 0.003 | 0.03 |
| bond angle (°) | 0.536 | 0.490 |
| Validation | |  |
| MolProbity Score | 1.61 | 1.15 |
| Clash score | 5.29 | 3.11 |
| Poor rotamers (%) | 0 | 0 |
| Ramachandran plot (%) |  |  |
| Favored | 98.13 | 97.78 |
| Allowed | 1.87 | 2.22 |
| Disallowed | 0 | 0 |

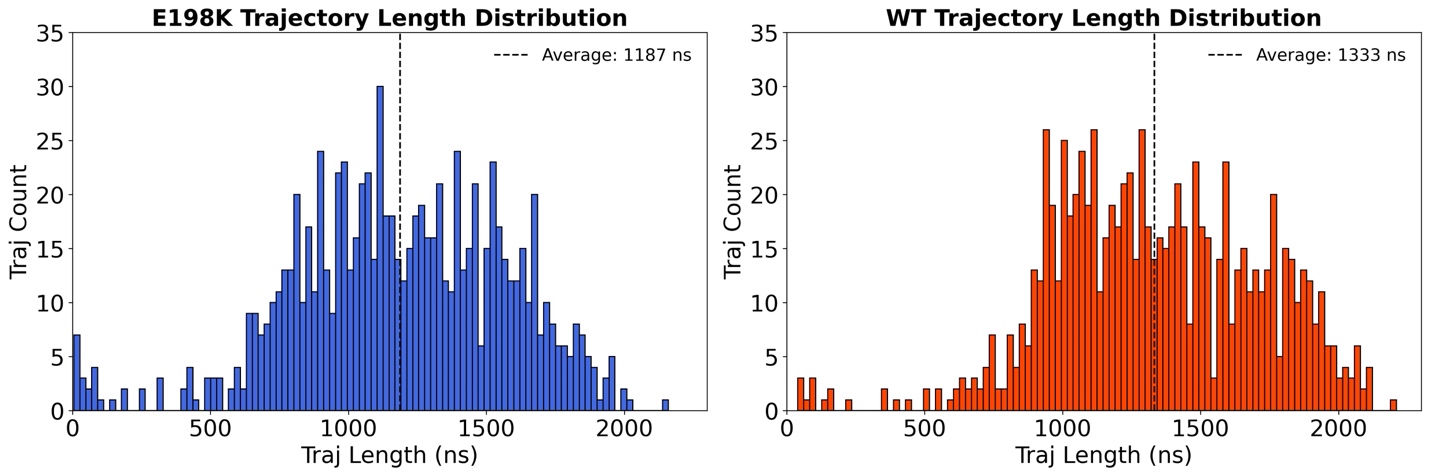
**Figure S2.** Histograms of unbiased MD trajectory lengths (ns) for E198K (left) and WT (right) collected using Folding@Home. The average lengths were 1,187 ns for E198K (1.04 ms total) and 1,333 ns for WT (1.17 ms total).

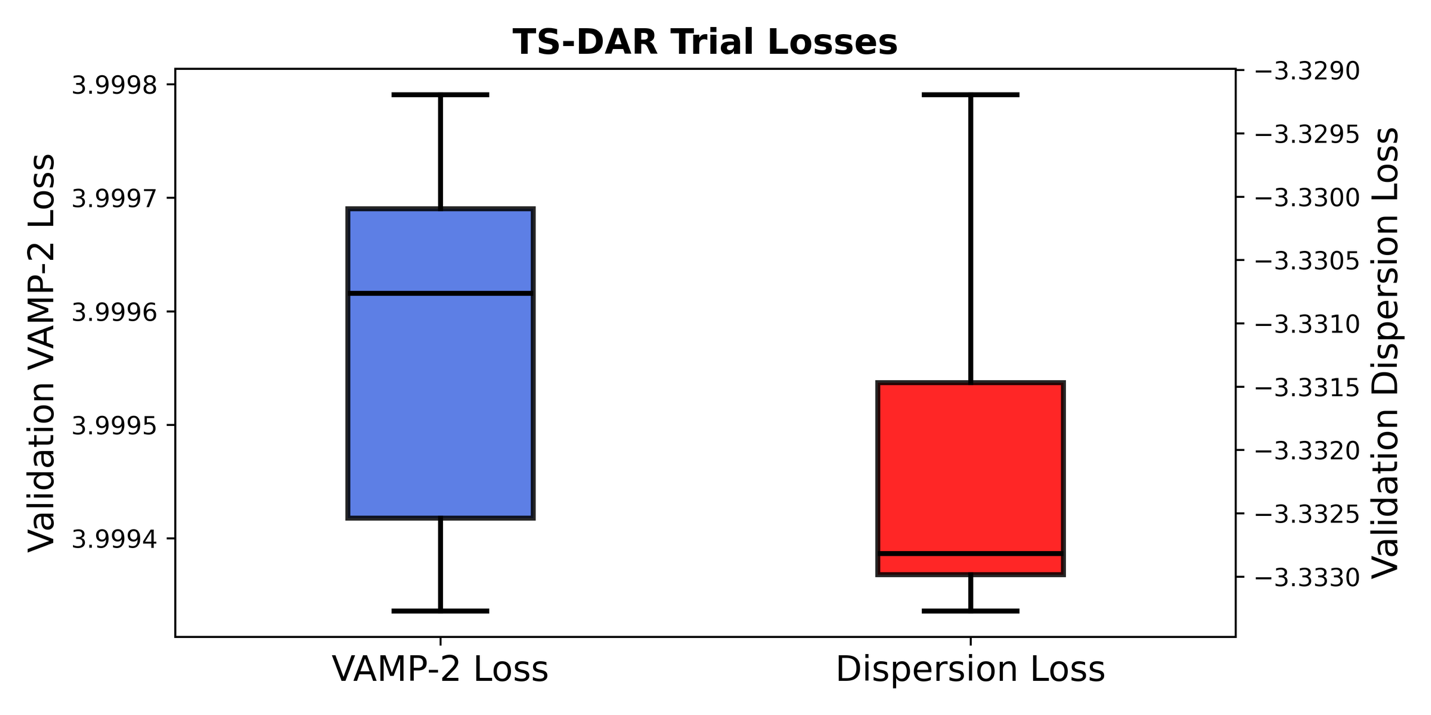

**Figure S3.** Box plot distribution of TS-DAR validation VAMP-2 and dispersion losses across 10 independent trained TS-DAR models starting from different initial seeds. The narrow distributions indicate that the model training was stable across independent runs and the learned kinetics were reproducible.

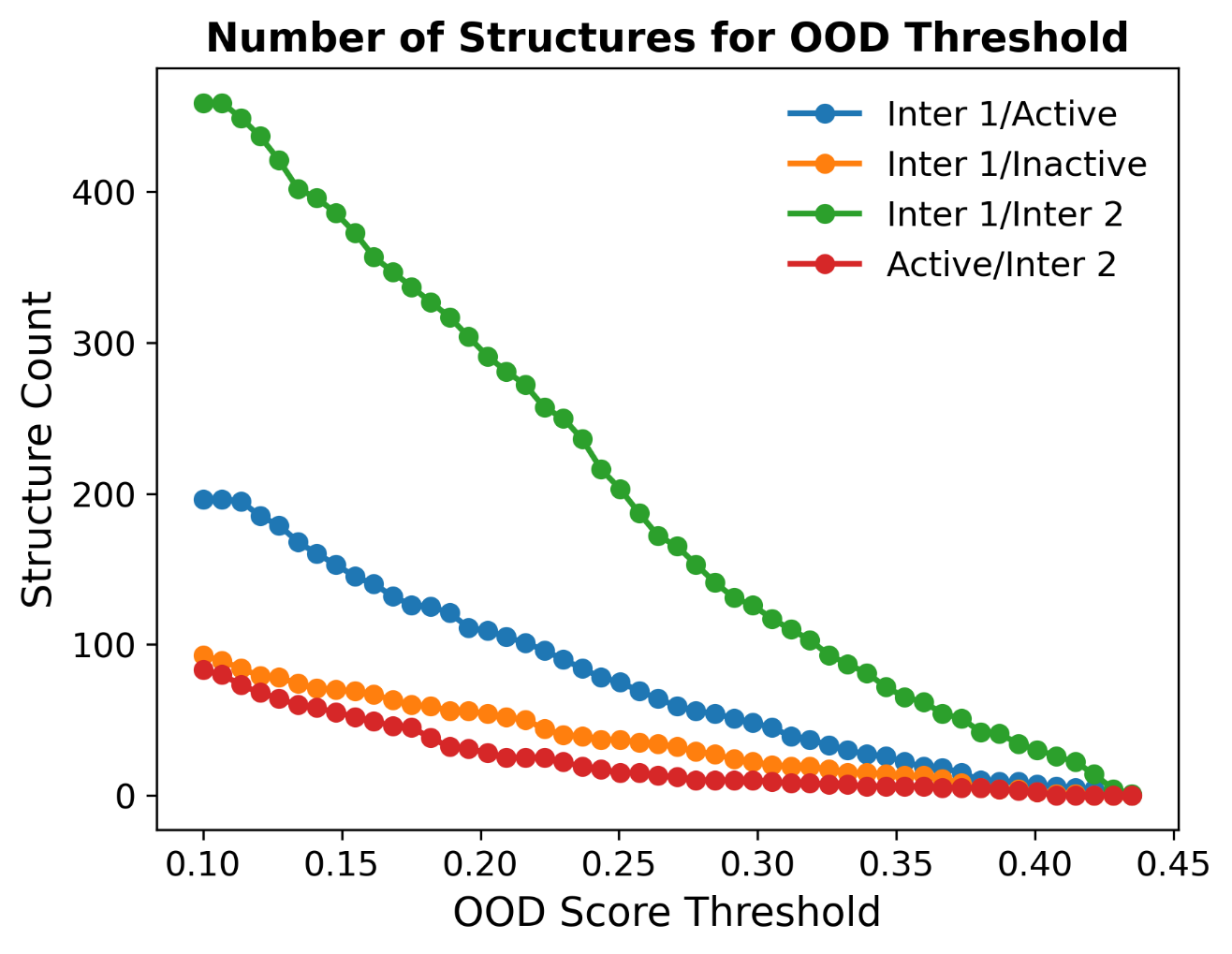

**Figure S4.** The number of E198K conformations assigned to each transition-state ensemble for pairs of macrostates were calculated at different OOD-score thresholds from the final TS-DAR model trained on the aggregate Folding@Home trajectories. As the OOD threshold increases, fewer conformations are retained.

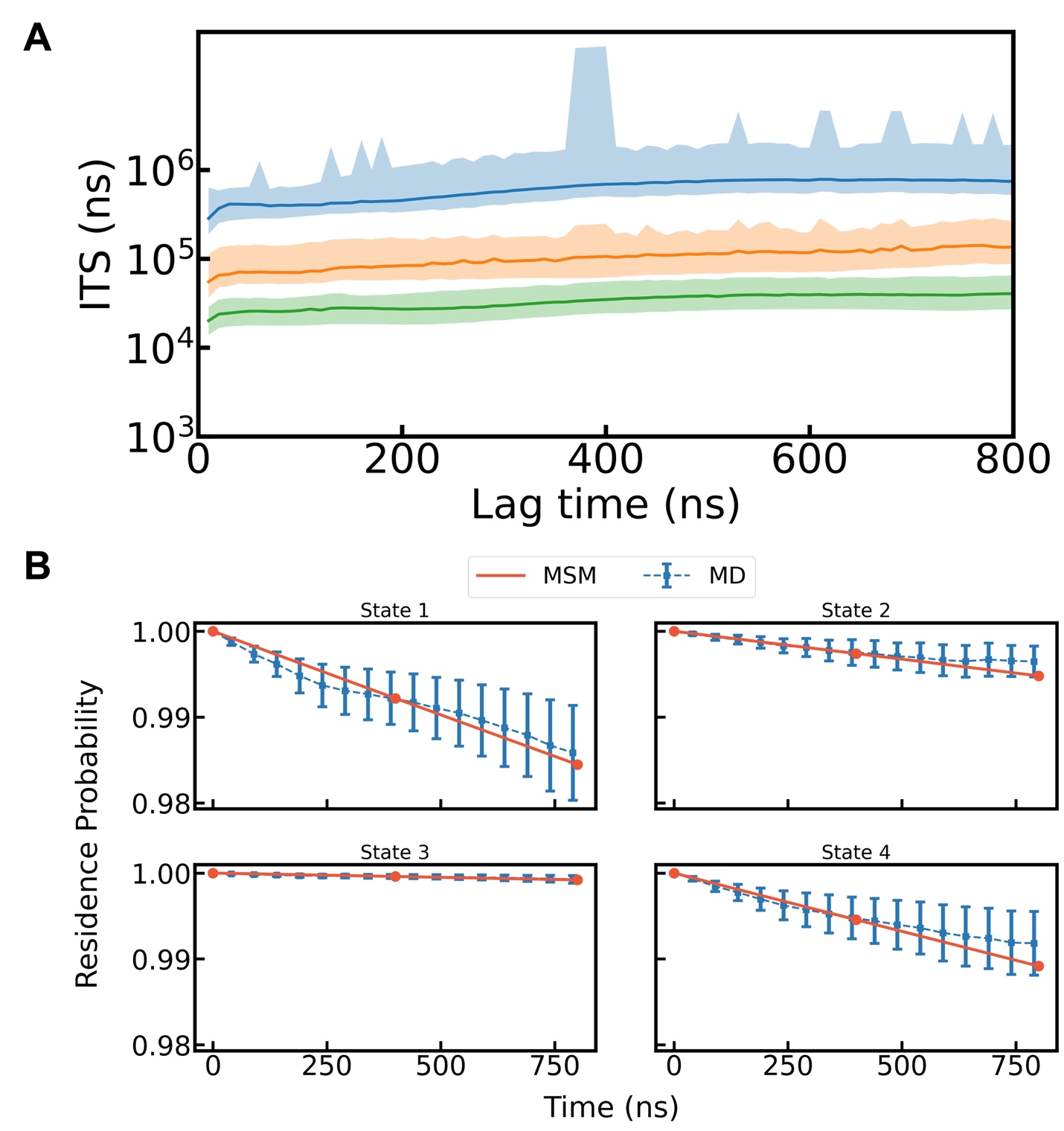

**Figure S5.** Validation of the four-state E198K MSM by Implied Time Scale (ITS) analysis and Chapman-Kolmogorov (CK) test. **(A)** ITS of the four-state E198K MSM as a function of lag time, where state assignments are obtained from final TS-DAR model. A lag time of 400 ns was chosen for the final MSM based on the plateau of the longest ITS. **(B)** CK test for the four-state E198K MSM comparing residence probabilities predicted by an MSM built at 400-ns lag time with those estimated directly from the unbiased MD trajectories. The error bars are calculated by performing 50 rounds of bootstrapping on the MD trajectories with replacement.

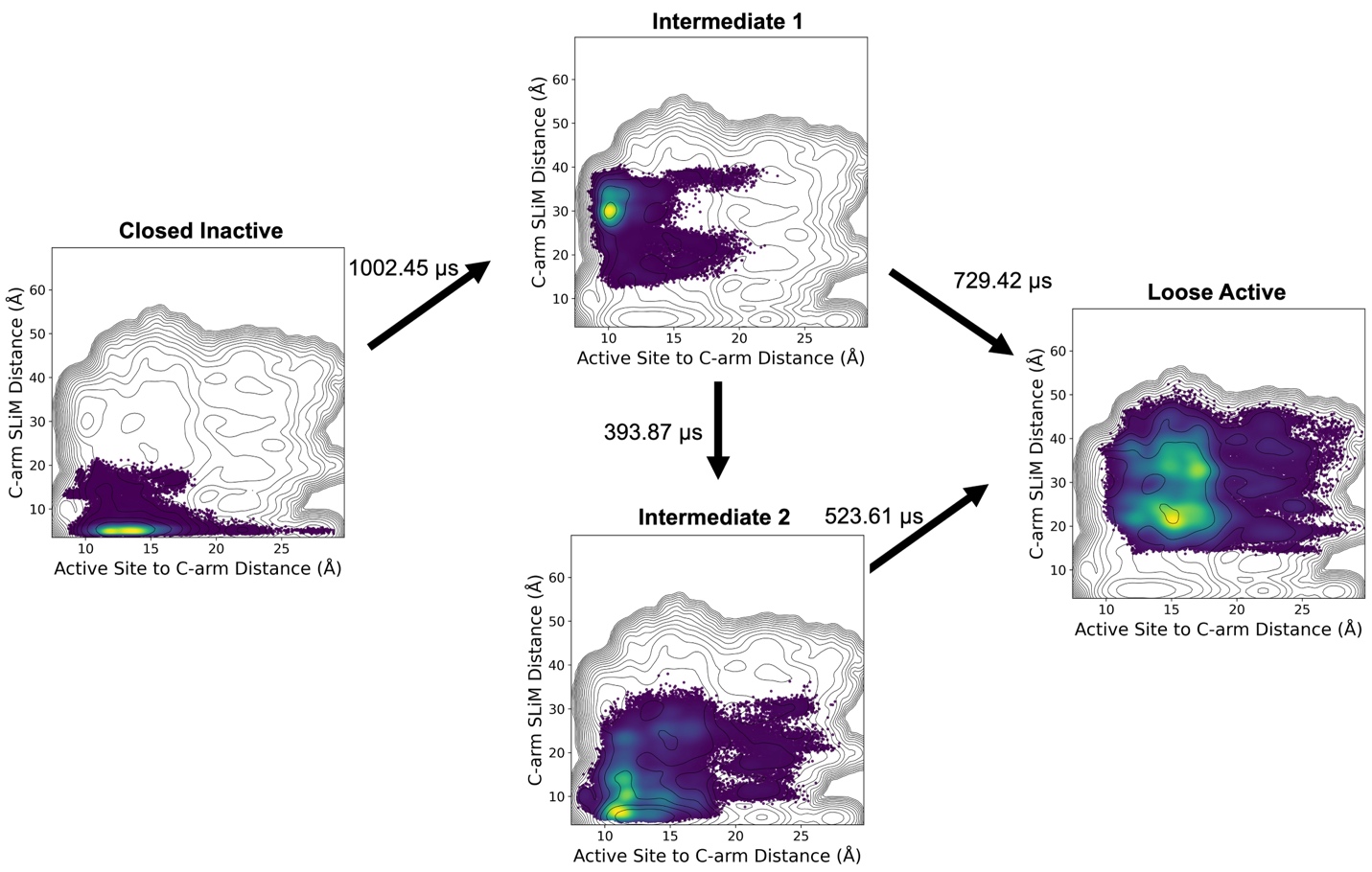

**Figure S6.** Four macrostates identified by TS-DAR projected onto the two-dimensional free energy surface for the “C-arm SLiM” distance (y-axis) and the “Active Site to C-arm” distance (x-axis) calculated from all the unbiased E198K MD simulations. The coloring is the density of frames calculated using kernel density estimate. The MFPTs are shown for each macrostate transition, calculated from the validated 400-ns lag time MSM.

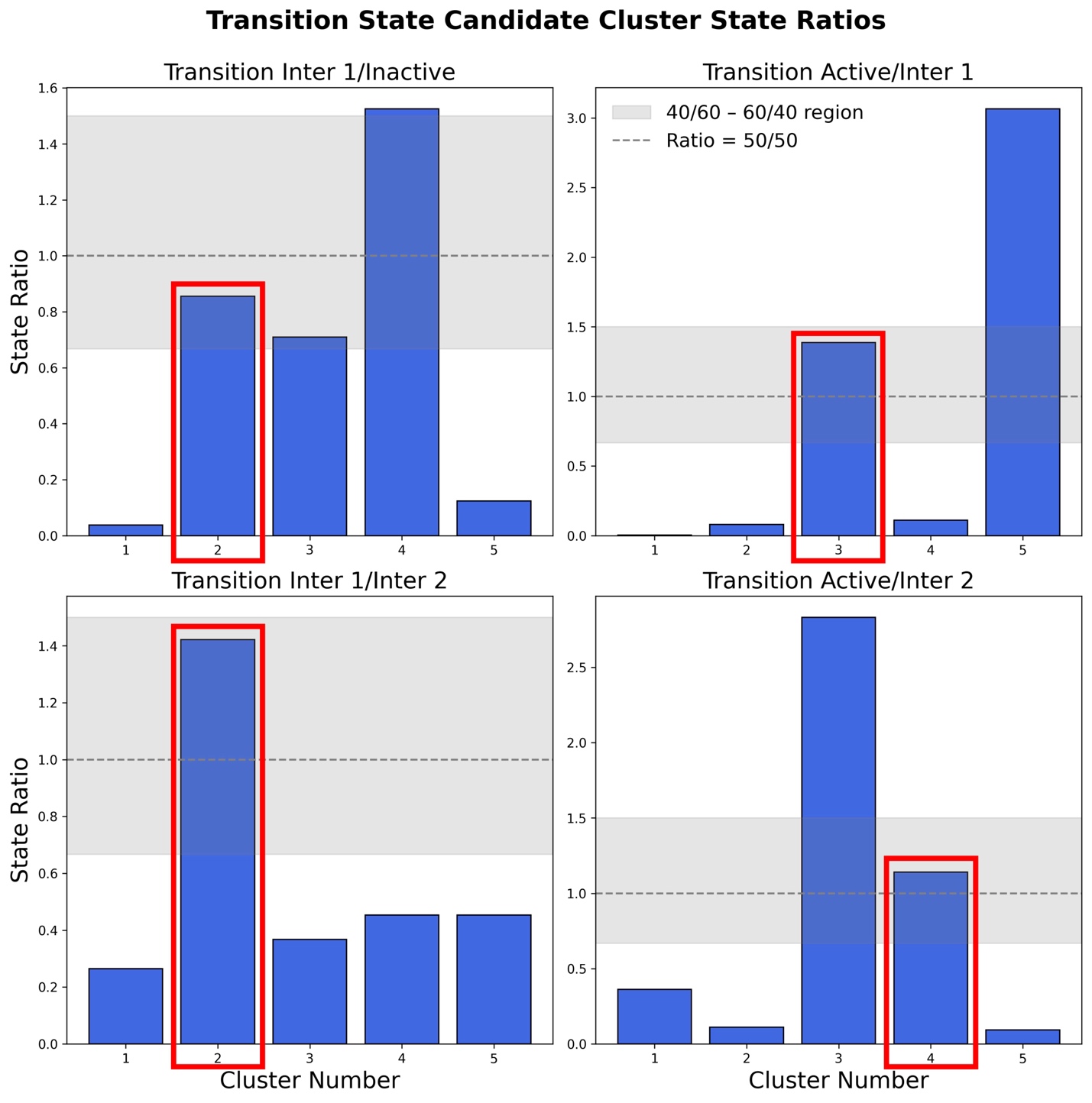

**Figure S7.** State ratios for simulations initiated from transition-state candidate structures. Candidate structures were chosen by clustering top 100 OOD scores for each transition identified by the E198K TS-DAR model (see Methods Section J in main text for details). The structures with the ratio of adjacent macrostates closest to one (50/50) were chosen to run WT perturbation simulations (highlighted with a red box).

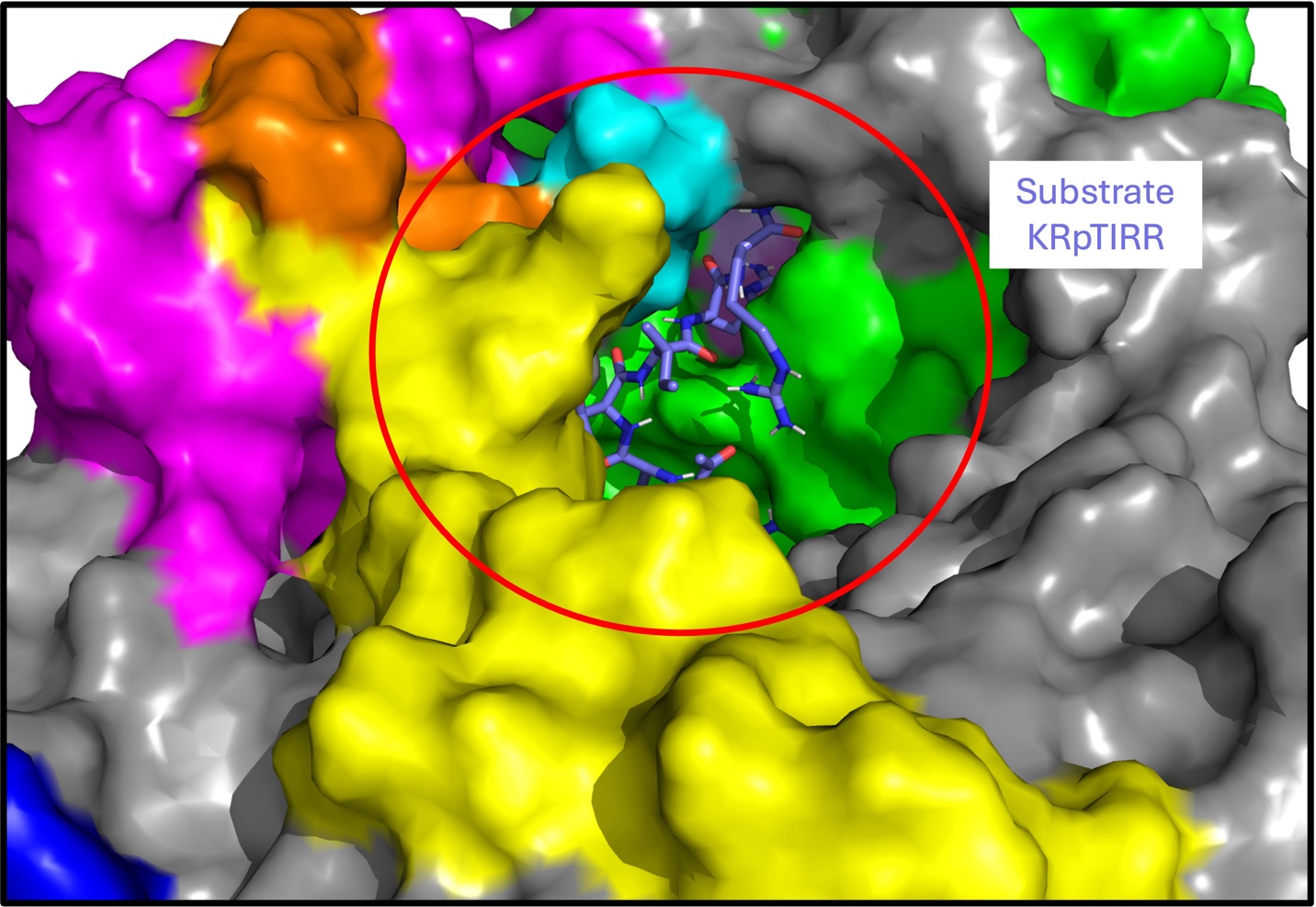

**Figure S8.** Docked structure of peptide substrate into E198K loose active state, identified by TS-DAR, showing direct access to the active site and space to accommodate the phosphorylated substrate. The docked hexapeptide (KRpTIRR) substrate was used for the *in vitro* phosphatase assay to determine the k_cat_ E198K and WT, taken from Konovalov et al. 2023, reproduced in Figure 4B. AutoDock Vina was used for the docking calculation.

**Table S2.** MFPTs for E198K macrostates.

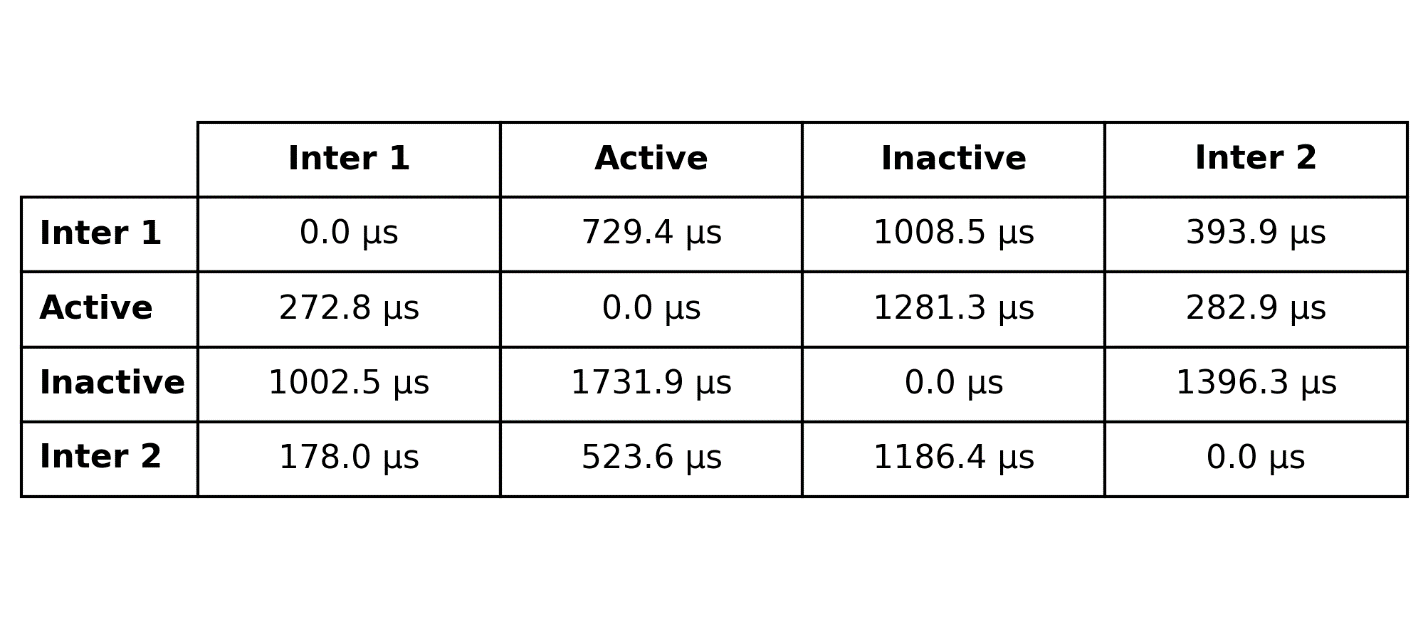
